# Single cell transcriptomic analysis of the adult mouse pituitary reveals a novel multi-hormone cell cluster and physiologic demand-induced lineage plasticity

**DOI:** 10.1101/475558

**Authors:** Yugong Ho, Peng Hu, Michael T. Peel, Sixing Chen, Pablo G. Camara, Douglas J. Epstein, Hao Wu, Stephen A. Liebhaber

## Abstract

The anterior pituitary gland drives a set of highly conserved physiologic processes in mammalian species. These hormonally-controlled processes are central to somatic growth, pubertal transformation, fertility, lactation, and metabolism. Current models, largely built upon candidate gene based immuno-histochemical and mRNA analyses, suggest that each of the seven hormones synthesized by the pituitary is produced by a specific and exclusive cell lineage. However, emerging evidence suggests more complex models of hormone specificity and cell plasticity. Here we have applied massively parallel single-cell RNA sequencing (scRNA-seq), in conjunction with a set of orthogonal mRNA and protein imaging studies, to systematically map the cellular composition of adult male and female mouse pituitaries at single-cell resolution and in the setting of major physiologic demands. These analyses reveal sex-specific cellular diversity associated with normal pituitary homeostasis, and identify an array of cells with complex complements of hormone-enrichment as well as a series of non-hormone producing interstitial and supporting cell lineages. These scRNA-seq studies identify a major cell population that is characterized by a unique multi-hormone gene expression profile. The detection of dynamic shifts in cellular representations and transcriptome profiles in response to two well-defined physiologic stresses suggests corresponding roles of a number of these clusters in cellular plasticity within the adult pituitary. These studies point to an unanticipated complexity and plasticity in pituitary cellular composition that expands upon current models and concepts of pituitary gene expression and hormone production.

## Introduction

The pituitary is a key regulatory gland in mammalian organisms. Complex arrays of hormonal outputs from the pituitary play central roles in physiological pathways and a wide array of inherited and acquired pathologic processes (Kelberman et al., 2009). These pathways impact post-natal growth, puberty, fertility, lactation, and metabolism. The pituitary contains a posterior lobe that comprises a direct extension of the central nervous system, and an anterior/median lobe (referred herein as anterior lobe) that is derived from oral ectoderm (Davis et al., 2013). The anterior pituitary contains cells that synthesize seven hormones: growth hormone (GH), prolactin (PRL), the β subunits of thyroid-stimulating hormone (TSHβ), luteinizing hormone β-subunit, (LHβ), and follicle-stimulating hormone β-subunit (FSHβ), as well as adrenocorticotrophic hormone (ACTH), and α melanocyte-stimulating hormone (α-MSH) (Fig. S1) (Zhu et al., 2007). Hormone synthesis and release from the anterior pituitary is coordinated by regulatory factors generated within signaling centers in the hypothalamus and transmitted to the pituitary *via* a dedicated hypothalamic-pituitary portal circulatory system (Vazquez-Borrego et al., 2018). A complete understanding of how these regulatory networks impact physiologic function requires defining the composition and relationships of cell lineages of the anterior pituitary and their corresponding patterns of hormone gene expression in an unbiased and comprehensive manner.

The prevailing model of anterior pituitary structure and function is based predominantly on the assumption that each hormone is expressed from a distinct cell type (Davis et al., 2013; Zhu et al., 2007); i.e. seven hormones secreted synthesized by a set of six corresponding cell lineages (in this model the two gonadotrope hormones, FSHβ and LHβ, are co-expressed from a single lineage) (Fig. S1). While compelling in many respects, this model is primarily based on targeted immuno-histochemical and mRNA analyses. The varying sensitivities and specificities of these approaches (Nakane, 1970), their inductive nature, and their limited capacity to examine multiple hormonal expressions within single cells, suggest that they may not be unbiased and fully informative. The possibility of a more complex composition of pituitary cell lineages and hormone expression has been suggested by previous reports of pituitary cells that express multiple hormones (Seuntjens et al., 2002a; Villalobos et al., 2004a, b). In addition, reports of pituitary cells with mitotic markers and factors linked to stem cell functions have been used to support complex models of pituitary dynamics linked to cellular expansion, *trans*-differentiation, and lineage plasticity. Thus, the degree to which the current model of hormone expression fully encompasses pituitary structure and function remains open to further study. This goal may be facilitated by assignment and cataloging of the gene expression profiles of individual cells in the pituitary using a comprehensive and unbiased approach.

The advent of single cell technologies has facilitated the analysis of cell lineages within a rapidly expanding array of tissues and has been used to effectively explore how defined cell lineages in various tissues respond to physiologic stresses and mediate specific functions (Tanay and Regev, 2017). Newly developed approaches of single cell isolation linked to high-throughput RNA sequencing approaches, such as Drop-Seq and 10X Genomics platforms (Macosko et al., 2015; Zheng et al., 2017), can now be applied to massively parallel analysis of single cell transcriptomes. These comprehensive and unbiased approaches can be of particular value in uncovering cellular heterogeneity and novel cell types, and in revealing corresponding regulatory factors involved in lineage differentiation and function (Campbell et al., 2017; Chen et al., 2017; Hu et al., 2017; Shekhar et al., 2016).

In the current study, we have applied Drop-seq technology (Macosko et al., 2015), in conjunction with a set of orthogonal single cell protein and RNA-based validation approaches, to define the steady state cellular composition of the adult mouse anterior pituitary at single cell resolution. Our data are concordant with certain aspects of current models, most notably in the identification of individual cell clusters expressing specific pituitary hormones and the presence of sexual dimorphism in pituitary cell compositions. Remarkably, these data also reveal the presence of a major cluster of multi-hormone expressing cells that contribute to the response of the pituitary to robust physiologic stresses linked to post-partum lactation and to stimulation by a hypothalamic regulatory factor. These analyses provide a comprehensive view of pituitary gene expression in adult pituitary and generate testable models of cell plasticity that underlie the capacity of the pituitary to respond to major physiologic demands.

## Results

### scRNA-seq analysis reveals an unexpected complexity of hormone gene expression within the *Pou1f1*^+^ clusters

Studies of pituitary development and lineage differentiation have suggested a model in which each of six distinct hormone expressing cell lineages expresses a corresponding polypeptide hormone (Fig. S1) (Zhu et al., 2007). The differentiation of these cell is controlled by a variety of transcription factors and signaling molecules (Kelberman et al., 2009). Multiple lines of genetic and biochemical evidence have converged on the conclusion that pituitary specific POU homeodomain transcription factor, Pou1f1, serves an essential function in driving terminal differentiation of cells expressing *Gh* (somatotropes), *Prl* (lactotropes), and *Tshb* (thyrotrope) (‘Pou1f1 dependent lineages’; Fig. S1) (Camper et al., 1990; Li et al., 1990). The most compelling support for this function is the observation that loss of *Pou1f1* gene expression results in the combined loss of *Gh, Prl*, and *Tshb* gene expression in both mice (Camper et al., 1990; Li et al., 1990) and humans (Ohta et al., 1992; Radovick et al., 1992). Based on these data, we performed single-cell transcriptomic analysis with the assumption that the Pou1f1 lineages would be organized into three discrete clusters, defined by their mutually exclusive expression of *Gh, Prl,* and *Tshb* mRNAs.

We employed Drop-seq technology to assess this model of *Pou1f1* lineage organization in an unbiased manner and to explore the full spectrum of pituitary cell composition (Macosko et al., 2015). Single cell transcriptome data were generated from 9 independent analyses including pituitaries harvested from different genders, ages, and physiologic conditions (Fig. S2). After excluding cells of low sequencing complexity (see **Methods** and Fig. S2), the transcriptomes of 18,797 cells were retained for downstream analysis. Dimensionality reduction and clustering of the full set of 18,797 single cell transcriptomes was based on principal component analysis (Macosko et al., 2015). Visualization of the data by Uniform Manifold Approximation and Projection (UMAP) (Stuart et al., 2019a), revealed nine spatially distinct clusters surrounding a set of three major clusters (Fig. S2).

We first analyzed the cellular clusters in the 7 to 8-week old, sexually naïve female and male mice (Fig. 1). Among the three major *Pou1f1*-expressing cell clusters identified in the UMAP plot (centrally located **in** Fig. 1B), the largest of them was assigned as somatotropes (Som) based on the high level enrichment for both *Gh* and the cell surface receptor for the hypothalamic growth hormone releasing hormone (*Ghrhr*) (see violin and feature plots Figs. 1B and C) (Lin et al., 1993). The second *Pou1f1*^+^ cluster was identified as lactotropes (Lac) based on the high level expression of *Prl* mRNA (Fig. 1C) in conjunction with markers previously linked to *Prl* expression and lactotrope functions; glutamine receptors (*Gria2*, *Grik2*, and *Grik5*) involved in PRL hormone release (Durand et al., 2008), two transcriptional co-activators of estrogen receptor function (*Ddx5* and *Ddx17*) (Janknecht, 2010), and two transcription factors recently identified as enriched in *Prl*^+^ cells and implicated in *Prl* gene activation (*Nr4a1* and *Nr4a2)* (**Table S1**) (Peel et al., 2018).

**Figure 1.**
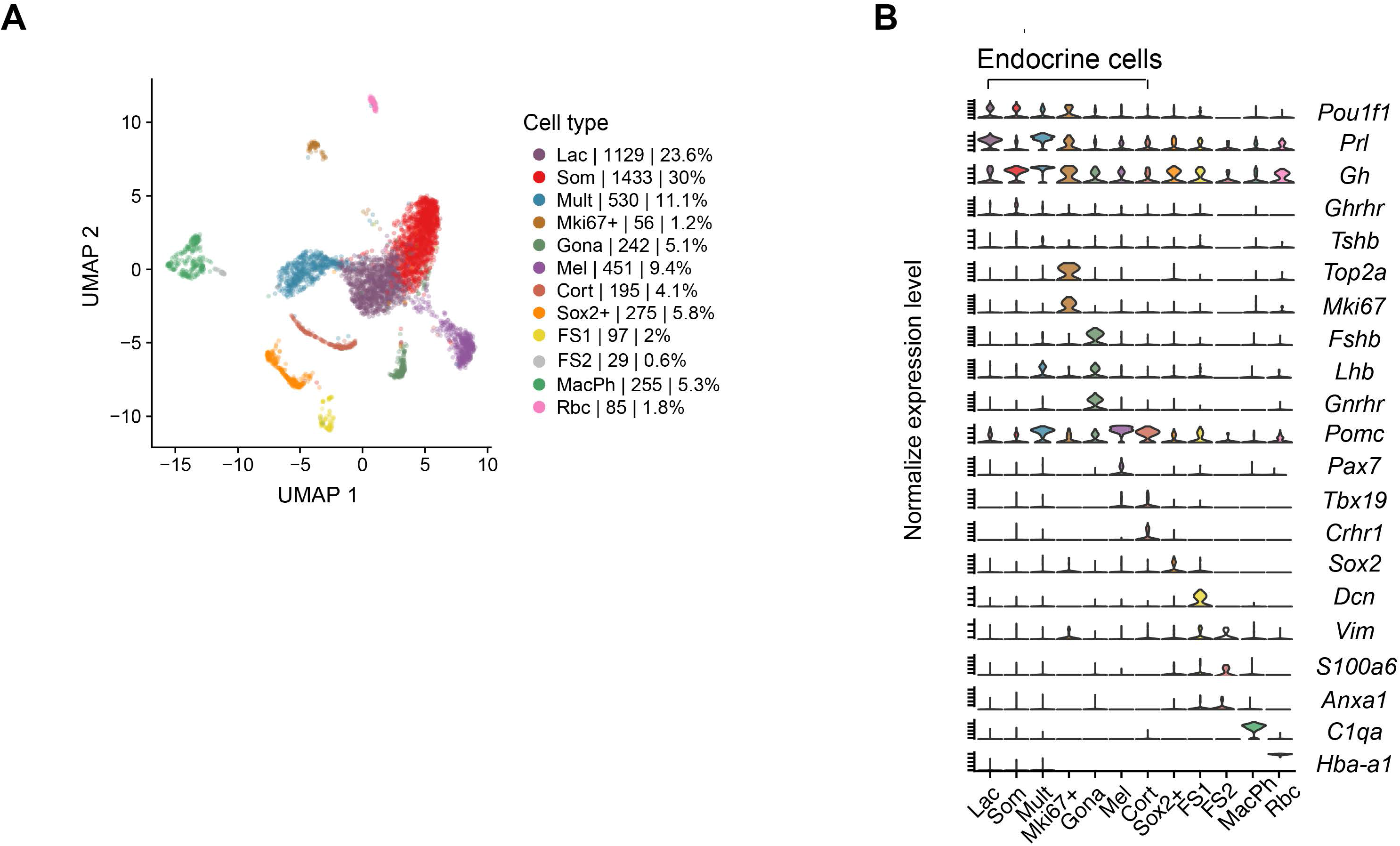
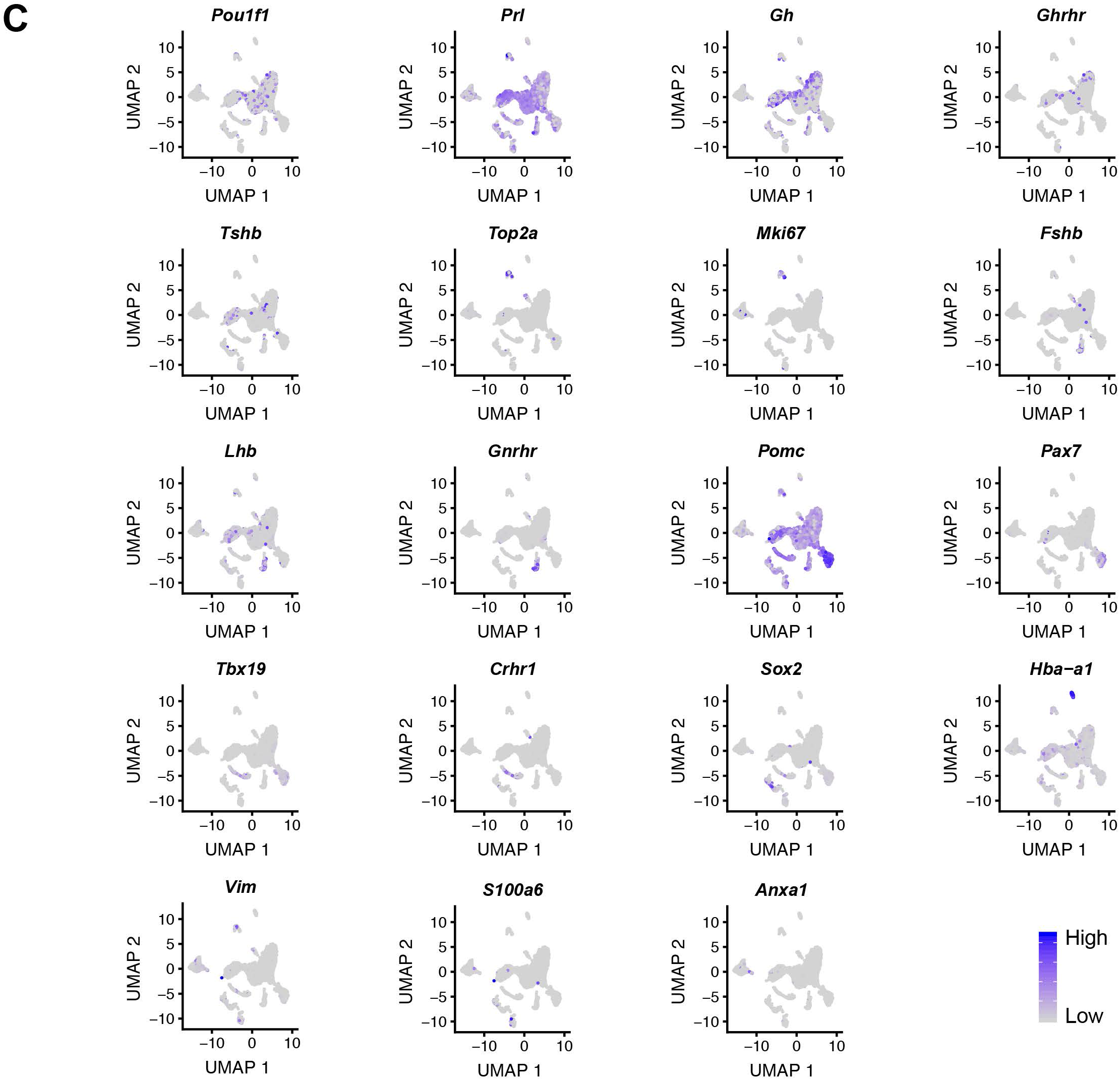
Single cell transcriptome analysis of the adult mouse pituitary. **A. Clustering of adult pituitary cells based on principal component analysis of single cell transcriptomes.** The spectral UMAP plot reflects the clustering analysis of individual adult pituitary cell transcriptomes (n = 4,777). The data represent the summation of 2 independent studies of pituitaries isolated from 7-8 week old, sexually naïve male and female mice. Each color-coded cluster was assigned an ‘identity’ based on the expression of sets of informative markers (as in **C** below; see text for details). The number and the percentage of cells in each cluster are indicated to the right of the UMAP plot. **B. Expression of a subset of the marker genes used to assign cell cluster identities.** The violin plots summarize mRNA expression corresponding to each indicated gene. The provisional identification of each cluster is noted at the bottom of each corresponding lane. Clusters corresponding to hormone expressing cells, including the *Mki67*+ cluster, are bracketed above the corresponding lanes. **C. Feature plots demonstrating the distributions of mRNAs encoding the pituitary transcription factors (*Pou1f1*, *Tbx19*, and *Pax7*), the signal receptors (*Ghrhr*, *Gnrhr*, and *Crhr1*), marker genes for each cluster, and six pituitary hormones.** These distributions are superimposed on the cell cluster diagram (**A**). Each feature map is labeled according to its respective mRNA. The heat map legend is displayed at the bottom right corner.

The characteristics of a third major *Pou1f1*^+^ cluster could not be clearly linked to any of the currently defined pituitary cell lineages. This Pou1f1-enriched cluster (‘Mult’, as defined below) encompassed 11.1 % of pituitary cells from 7 to 8-week old CD1 mice (Fig. 1A) and was remarkable in several respects. First, the co-expression of *Gh* and *Prl* mRNAs was detected at high levels that were comparable to those detected in the somatotrope (Som) and lactotrope (Lac) clusters (Fig. 1B). Secondly, the cells in this cluster were also highly enriched for mRNAs transcribed from the *Pomc* gene at levels comparable to that detected in the ‘Pou1f1-independent’ melanotrope and corticotrope lineages (Fig. 1B **and** C) (Zhu et al., 2007). Thirdly, substantial number of cells in this third cluster also contained mRNAs encoding the gonadotrope hormone, *Lhb* (Fig. 1B and C). Lastly, 32% of the cells in the anterior pituitary that contained *Tshb* mRNA mapped to this cluster (Fig. 1C). On the basis of these observations, we provisionally assigned this cluster as a ‘multi-hormone’ producing cluster (Mult). Further analysis of the differentially expressed genes revealed that several genes involved in metabolism (*Gpx3*, *Ddah1*, and *Nme1*) were exclusively enriched in this cluster (**Table S1**). In summary these data revealed a novel *Pou1f1*^+^ multi-hormone producing cluster that comprises a substantial fraction of the pituitary cell population.

The scRNA seq analysis revealed a fourth *Pou1f1*^+^cell cluster, substantially smaller than the three noted above, that expresses proliferating cell markers such as *Top2a* and *Mki67* (Fig. 1B). This observation is consistent with prior reports that the majority of proliferating cells in the adult pituitary are positive for *Pou1f1* expression (Cao et al., 2016; Zhu et al., 2015).

In summary the single cell analysis of the adult pituitary revealed three major *Pou1f1*^+^ cell clusters representing the expected populations of somatotropes and lactotropes, predominantly expressing *Gh* and *Prl* mRNAs, respectively, as well as a novel *Pou1f1*^+^ cluster containing cells with a complex pattern of multi-hormone gene expression.

### Single cell RNA-seq identifies POU1F1 independent hormone-producing cell lineages and non-hormonal cell clusters in the adult pituitary

The eight remaining cell clusters that identified in the UMAP plot lacked the high level of *Pou1f1* mRNA (Fig. 1B **and** C). Three of these eight clusters could be directly assigned to specific hormone-expressing cell lineages based on patterns of marker gene expression (Fig. 1B **and** C). A cluster corresponding to melanotropes (‘Mel’) was assigned based on the co-expression of pro-opiomelanocortin (*Pomc)* prohormone mRNA, the transcription factor *Tbx19*, and the melanotrope-restricted paired homeodomain transcription factor *Pax7* (Budry et al., 2012; Mayran et al., 2018). A second cluster was assigned as a corticotrope cell cluster (‘Cort’). These cells shared with the melanotrope cluster in enrichment for *Pomc* and *Tbx19* mRNAs but lacked substantial level of *Pax7* mRNA (Lamolet et al., 2001; Liu et al., 2006; Philips et al., 1997). A third *Pou1f1*-negative cluster was assigned to the gonadotrope lineage (‘Gona’ cluster) based on the enrichment for mRNAs encoding the two gonadotrope-specific hormone subunits, *Fshb* and *Lhb*, in conjunction with the gonadotrope-restricted cell surface receptor for the hypothalamic regulatory factor, *Gnrh* (Ingraham et al., 1994; Zhu et al., 2007). Thus the initial clustering analysis of the single cell transcriptomes from the adult mouse pituitary identified a set of *Pou1f1*-enriched clusters and a set of clusters corresponding to three *Pou1f1*-independent lineages (melanotropes, corticotropes, and gonadotropes) (Fig. 1A).

The scRNA-seq data sets identified five additional clusters that represented non-hormonal cell lineages. Two of these clusters were identified as folliculostellate (FS) cells based on known marker genes (‘FS1 and 2’; Fig. 1A **and** B). FS cells have been proposed to serve a variety of structural, paracrine, and/or support functions in the pituitary (Le Tissier et al., 2012). Histologic identification of these cells has been based on an array of protein markers including *Anxa1*, *Vim*, and *S100* (Devnath and Inoue, 2008; Nakajima et al., 1980; Theogaraj et al., 2005). Differences in the relative expression levels of these markers has been used to define sub-types of FS cells in the anterior pituitary (Allaerts and Vankelecom, 2005).

The largest cluster among the non-hormonal lineages, representing 5.8% of the cell in the analysis, was assigned as a putative stem cell cluster (‘Sox2^+’^ clusters; Fig. 1A). This assignment was based on the shared expression of the progenitor/stem marker *Sox2* (Fig. 1B) (Andoniadou et al., 2013; Rizzoti et al., 2013). A separate cluster was identified as macrophage (‘MacPh’ cluster) based on the expression of the known macrophage marker gene, *C1qa* (Zahuczky et al., 2011) (Fig 1B). An erythrocyte cell cluster, defined by robust globin gene expression (Rbc cluster in Fig. 1) was assumed to originate from blood in the dissected pituitaries. In summary, these data reveal four cell clusters in the pituitary that serve functions other than direct production of polypeptide hormones; two FS cell clusters, a stem cell cluster, and a cluster representing a macrophage population.

To assess platform-specific bias in our study, we compared the Drop-seq analysis (above) with a transcriptome profile of 3,927 cells using 10X Genomic platform (cells isolated from four 7-8 week old, sexually naïve CD mice (Cheung et al., 2018; Zheng et al., 2017) (See methods and Fig. S3). This analysis using the 10X platform identified all of the cellular clusters identified using the Drop-seq platform (Fig. S3). Of particular note was the comparable identification of a multi-hormone cluster between Drop-seq (11.1%) and 10X Genomics (14.8%) data sets (Fig. S3). These results support the technical robustness and biological reproducibility of our scRNA-seq analysis.

In summary, the analysis of single cell transcriptomes from the adult mouse pituitary identifies eight hormone-expressing clusters, two clusters of folliculostellate cells, a set of putative stem cell clusters, and a single cluster of corresponding to a macrophage population.

### Single cell mRNA and protein analyses validate the prevalence of multi-hormone co-expression

The Drop-seq analysis revealed that 31.6% (995/3,148) of the cells in *Pou1f1*+ clusters of the 7 to 8-week old mice were positive for both *Gh* and *Prl* mRNAs (*Gh*+/*Prl*+). As an initial step in validating this prevalent co-expression of these two hormone mRNAs, we probed tissue sections and dissociated cells from adult male and female pituitaries by single cell RNA <u>f</u>luorescent in <u>s</u>itu <u>h</u>ybridization (scRNA FISH). Pituitary tissue sections of 8-week old mice were hybridized with arrays of fluorescent oligo probes antisense to the *Prl* and *Gh* mRNAs (Fig. 2 and S4A). As expected, *Prl* and *Gh* expression was observed to be restricted to the anterior lobe (AL) while *Pomc* expression was detected in both the intermediate lobe (IL) and AL (Fig. S4A). The RNA-FISH analysis further confirmed that a fraction of the *Gh* and *Prl* expressing cells were dual positive for these two mRNAs (Fig. 2A and B). The level of co-expression of *Gh* and *Prl* mRNAs was determined by analysis of 5 or more randomly selected fields (> 600 cells per pituitary) of pituitary tissue sections. This analysis revealed that 30.8% of all the *Gh* and/or *Prl* expressing cells in the female pituitary co-expressed *Prl* and *Gh* mRNAs (*Gh*^+^/*Prl*^+^) while 25.8% were uniquely positive for *Gh* mRNA (*Gh*^+^/*Prl*^-^) and 43.4% cells were uniquely positive for *Prl* mRNA (Fig. 2A**, right graph**). A parallel study of pituitary cells from male mice revealed 31% of the cells as dual positive (*Gh*^+^/*Prl*^+^), 52.3% as uniquely *Gh* positive (*Gh*^+^/*Prl*^-^), and 16.7% as uniquely *Prl* mRNA positive (*Prl*^+^/*Gh*^-^) (Fig. 2B**, right graph**). We conclude from these studies that co-expression *Gh* and *Prl* mRNAs is prevalent in the *Pou1f1* cell population in the adult pituitary. These studies also revealed a marked sexual dimorphism in the representation of cells that express *Gh* in the absence of *Pr*l and *Prl* in the absence of *Gh*.

**Figure 2.**
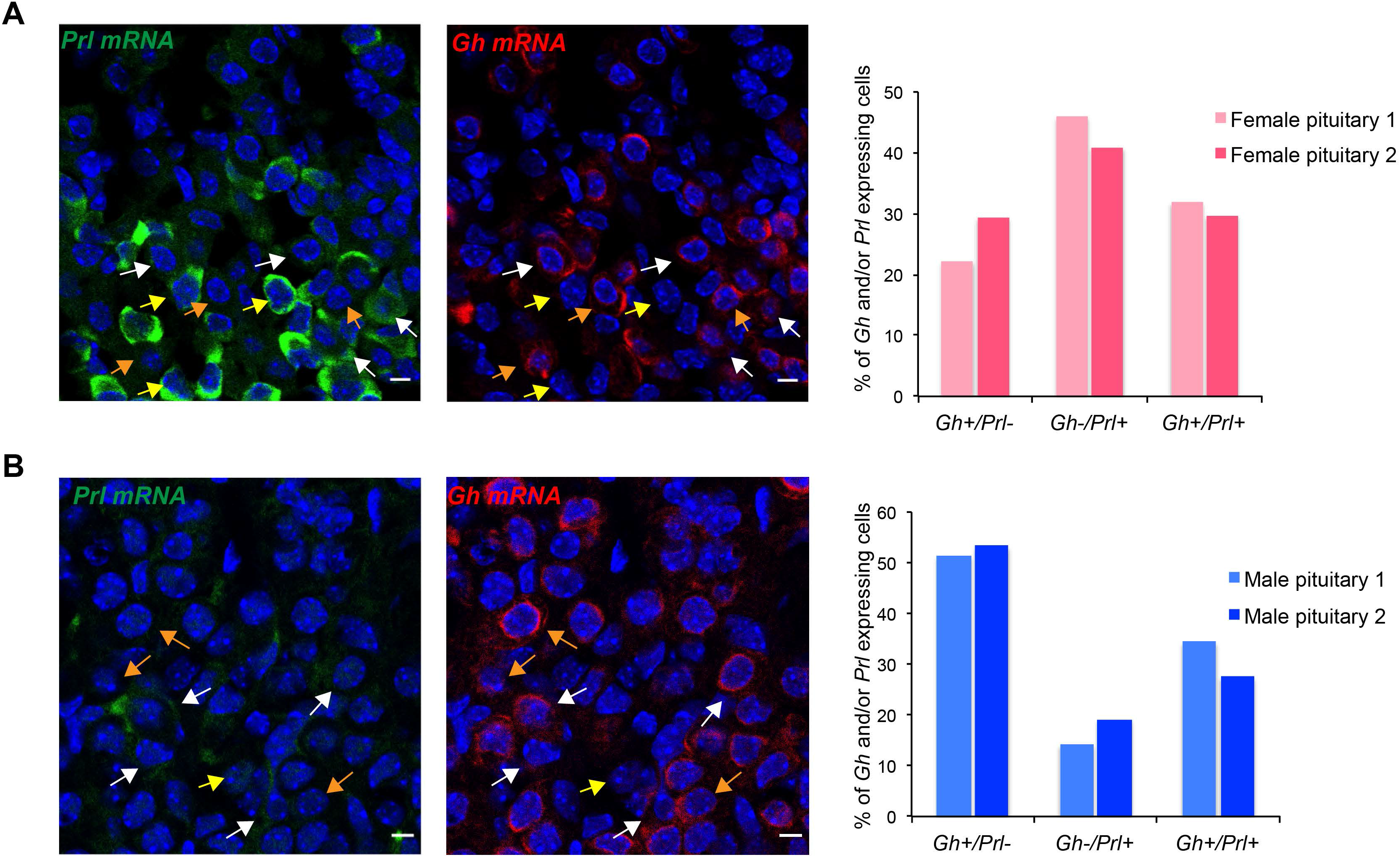
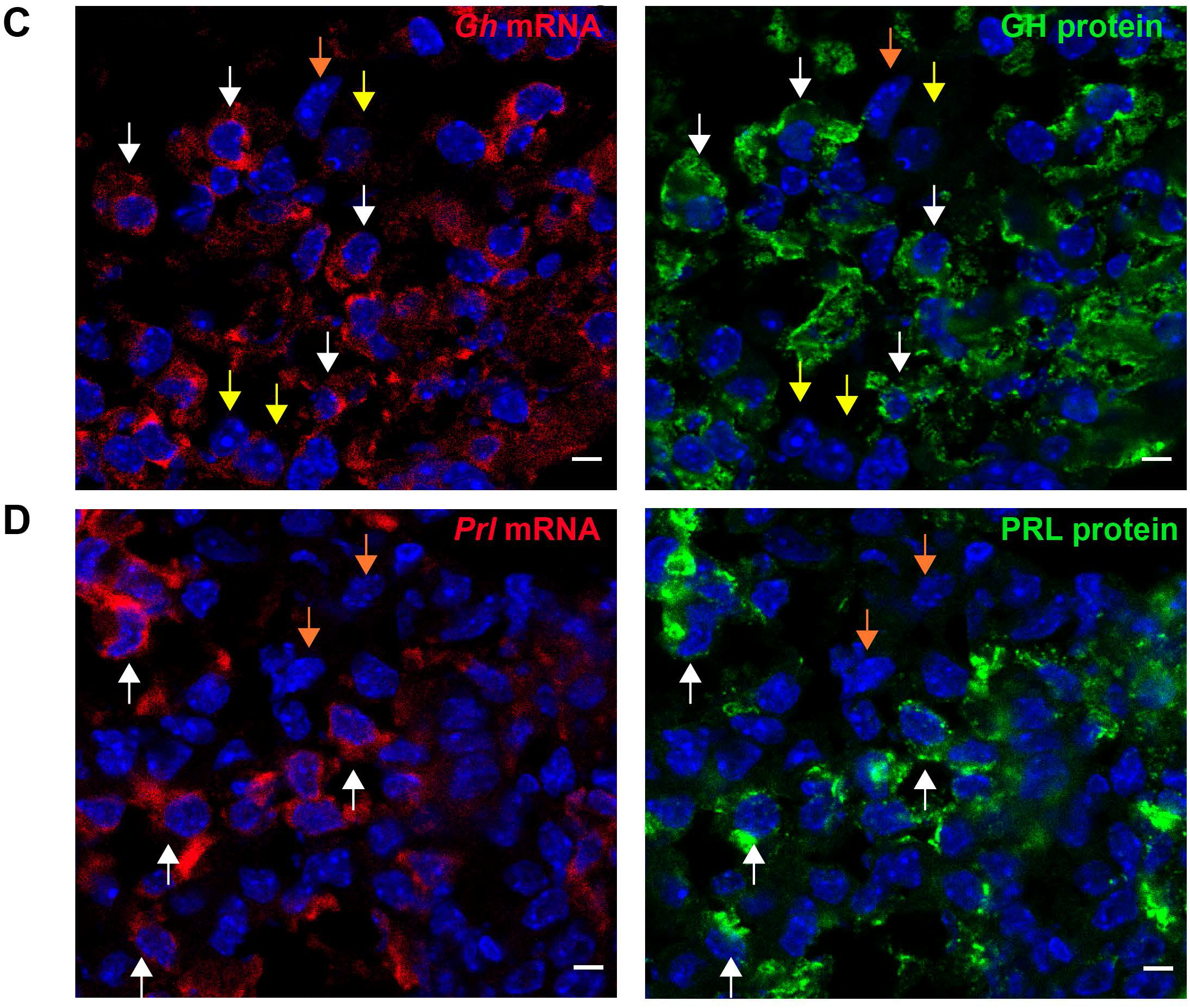
Single cell RNA fluorescent *in situ* hybridization (scRNA FISH) analysis detects the co-repression of *Gh* and *Prl* mRNAs in cells in the mouse pituitary. **A. ScRNA FISH analysis of female adult mouse pituitary tissue sections reveals a concordance of *Gh* with *Prl* expression *in vivo*.** **Left panel: *Prl* mRNA detection.** *Prl* mRNAs were detected by FISH using an array of anti-sense oligonucleotide probes (see **Methods)**. Bars = 5 µm. **Right panel: *Gh* mRNA detection**. *GH* mRNA was detected by FISH using an array of anti-sense oligonucleotide probes (see **Methods)**. Bars = 5 µm The white arrows in the two frames indicate four examples of *Gh* and *Prl* dual positive cells. The orange arrows indicate three cells with robust *Gh* mRNA expression but without *Prl* mRNA expression. The yellow arrows indicate three cells with robust *Prl* mRNA expression but without *Gh* mRNA expression. **Histogram: analysis of female pituitary cells for *Gh* and/or *Prl* mRNA. Pituitary tissue sections** from 8-week old, sexually naïve adult female mice (n=2) were analyzed by RNA FISH. Cells from 5 randomly selected images from each pituitary were analyzed (>600 cells per pituitary). The histogram displays the representation of cells positive for *Gh* but not *Prl* mRNA (*Gh*^+^*Prl*^-^; 25.8%), cells positive for *Prl* but not *Gh* mRNA(*Gh*^-^*Prl*^+^; 43.4%), and those dual positive cells (*Gh*^+^*Prl*^-^; 30.8%) in *Gh* and/or *Prl* expressing cells. **B. scRNA FISH analysis of male adult mouse pituitary tissue sections reveals a concordance of *Gh* with *Prl* expression *in vivo*.** **Left panel: *Prl* mRNA detection.** *Prl* mRNA was detected by FISH using an array of anti-sense oligonucleotide probes as described in **Methods**. Bars = 5 µm. **Right panel: *Gh* mRNA detection**. *GH* mRNA was detected by FISH using an array of anti-sense oligonucleotide probes as described in **Methods**. Bars = 5 µm The white arrows indicate four representatives of the *Gh* and *Prl* dual positive cells. The orange arrows indicate three cells with robust *Gh* mRNA expression but without *Prl* mRNA expression. The yellow arrow indicates a cell with *Prl* mRNA expression but without *Gh* mRNA expression. **Histogram: analysis of male pituitary cells. Pituitary tissue sections** from 8-week old, sexually naïve male mice (n=2) were analyzed by scRNA FISH for *Gh* and *Prl* mRNA expression as described in **A**. The histogram represents the percent of the total cell population positive for *Gh* but not *Prl* mRNA (*Gh*^+^*Prl*^-^; 52.3%), positive for *Prl* but not *Gh* mRNA (*Gh*^-^*Prl*^+^; 16.7%), and positive for both *Gh* and *Prl* mRNA (*Gh*^+^*Prl*^+^ 31%) in *Gh* and/or *Prl* expressing cells. **C. Combined single cell RNA FISH and IF analyses (scRNA FISH/IF) confirm the concordance of hormone mRNA and protein expressions.** **Top panels: Detection of *Gh* mRNA and GH protein.** The scRNA FISH analysis for *Gh* mRNA and IF analysis for GH protein were performed in the pituitary tissue sections of male adult pituitary (see **Methods)**. **Left panel: detection of *Gh* mRNA by scRNA FISH.** Scale bar=5 µm. **Right panel: detection of GH proteins by IF analysis in the some field**. Scale bar; 5 µm. The white arrows indicate the cells with robust signals for both *Gh* mRNA and GH protein. The yellow arrows indicated cells with trace levels of *Gh* mRNA and lack of GH protein signal. The orange arrow indicates a cell lacking *Gh* mRNA and GH protein signals. **Bottom panels: Correlation of *Prl* mRNA and PRL protein.** The scRNA FISH analysis of *Prl* mRNA and IF analysis of PRL protein were performed in female adult pituitaries (see **Methods)**. **Left panel: detection of *Prl* mRNA by scRNA FISH analysis.** Scale bar=5 µm. **Right panel: detection of GH protein by IF analysis in the same field.** Scale bar; 5 µm. The white arrows indicate four cells with robust signals for both *Prl* mRNA and PRL protein. The orange arrows indicate two cells lacking *Prl* mRNA and PRL protein signals.

To further explore the prevalent co-expression *Gh* and *Prl* genes, we next combined IF analyses of GH and PRL hormone proteins with scRNA FISH of the corresponding hormone mRNAs in pituitary tissue sections (Fig. 2C). Analysis of 933 cells confirmed concordance of mRNA and protein expression from these two genes; all cells with *Gh* or *Prl* mRNA as visualized by scRNA FISH had a comparable IF signal for the corresponding protein (Fig. 2C); when *Gh* or *Prl* mRNA was abundant the corresponding protein was also abundant and when either mRNA was at trace levels or undetectable the corresponding protein was comparably trace or negative by IF (examples in Fig. 2C). In no case did we observe a substantial discordance between corresponding IF and scRNA FISH signals.

We next assessed the prevalence of GH and PRL co-expression at the protein level (Fig. S4B). A double IF for GH and PRL analysis was performed in the tissue sections of an 8-week old male mouse and an 8-week old female mouse. We observed that a significant proportion of cells expressing either GH or PRL co-expressed both of the hormones (14% in the males and 19% in the females) (Fig. S4B). The difference in the percentages of dual positive cells detected by RNA-FISH and by IF (Figs. 2 and S4B) most likely reflects the different sensitivities of these two approaches as indicated in our combined RNA-FISH and IF studies (Fig. 2C).

*Pomc* mRNA encodes a prohormone that generates multiple smaller peptide hormones by site-specific cleavages specific to particular cells in the pituitary (Cawley et al., 2016). Two of these peptides, ACTH and α-MSH are lineage specific markers for anterior lobe (AL) corticotropes and intermediate lobe (IL) melanotropes, respectively (Zhu et al., 2007). The mapping of *Pomc* expression by Drop-seq revealed that *Pomc* mRNA was present at high levels (equivalent to that detected in the corticotrope lineage) in the cells that constituted the multi-hormone cluster (Fig. 1B **and** C) and was co-expressed with *Gh* and *Prl* mRNAs. This observation was validated by scRNA FISH. Combined RNA-FISH analyses of *Pomc* and *Prl* mRNAs revealed that majority of the *Prl* mRNA positive cells were also positive for *Pomc* mRNA (Fig. 3B). Remarkably, scRNA FISH analysis in female pituitary cells (n=3 pituitaries) revealed that 74% of the cells co-expressed *Pomc* and *Prl* and only 9.8% of the cells were *Pomc*^+^ but *Prl^-^* (Fig. 3C**, left graph**). A second scRNA FISH analysis in male pituitary cells (n= 3 pituitaries) revealed that 59% of all analyzed cells from male pituitary (n=3 pituitaries) co-expressed both *Pomc* and *Gh* while only 18.3% of the cells were *Pomc^+^* but *Gh*^-^ (Fig. 3B**, right graph**). These scRNA FISH studies are concordant with the initial Drop-seq data and lead us to conclude that the *Pomc* gene is co-expressed with the *Gh* and/or *Prl* gene(s) in majority of cells in the AL of the adult mouse pituitary.

**Figure 3.**
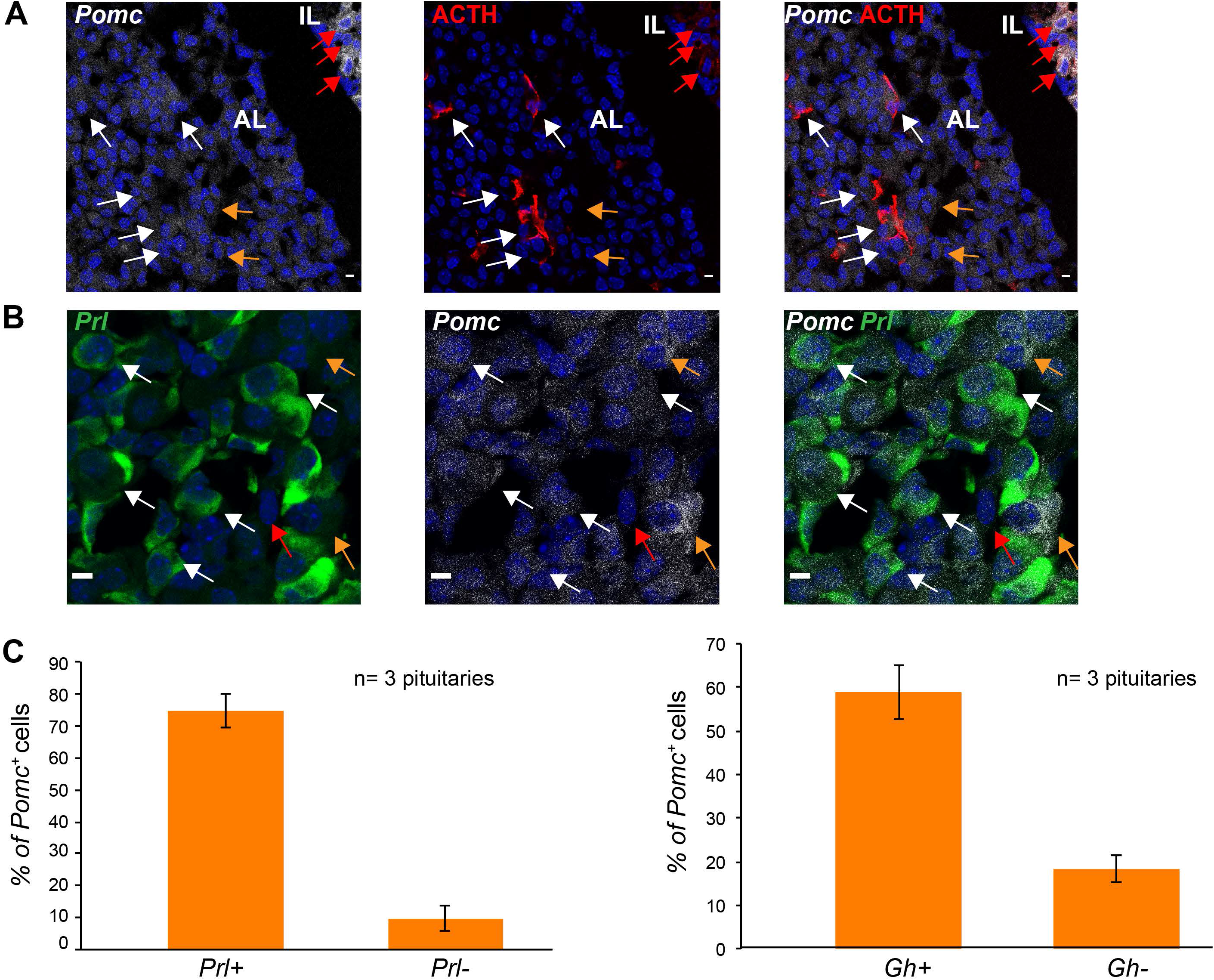
*Pomc* mRNA is co-expressed with the *Prl* mRNA and *Gh* mRNA in the anterior lobe of the adult pituitary. **A. *Pomc* mRNA is broadly expressed in the anterior lobe (AL) of the adult pituitary.** **Left panel:** *Pomc* mRNA (gray) were detected by RNA FISH analysis using anti-sense oligonucleotide probes (see **Methods)**. Scale bar = 5 µm. **Middle panel:** ACTH protein (red) was detected by IF analysis using antibody against ACTH. The ACTH antibody also recognizes α-MSH, the peptide hormone generated by the cleavage of ACTH in the intermediate lobe (IL). Scale bar = 5 µm **Right panel:** The merged images of the *Pomc* RNA FISH and ACTH IF in the same field. Scale bar = 5 µm, The white arrows indicate cells with *Pomc* mRNA and robust levels of ACTH protein in the AL. The orange arrows indicate cells with *Pomc* mRNA in non-corticotropes (ACTH negative) in AL. The merged images further confirm the robust expression of *Pomc* mRNA in the melanotropes (α-MSH+) (red arrows in the IL). **B. RNA FISH analysis detects co-expression of *Pomc* mRNA and *Prl* mRNA in cells of the anterior lobe of adult pituitary.** *Prl* mRNA was detected by FISH using an array of 488-conjugated anti-sense oligonucleotide probes (green in left panel). *Pomc* mRNA was detected by FISH using an array of Cy5-conjugated anti-sense oligonucleotide probes (gray in middle panel). The right panel is the merged images of the *Prl* mRNA FISH and *Pomc* RNA FISH in the same field. The white arrows indicate cells with co-expression of *Prl* and *Pomc* mRNAs, the orange arrows indicate cells with robust signal for *Pomc* mRNA and without *Prl* mRNA, and the red arrow indicates a cell that is negative for both *Prl* and *Pomc* mRNAs. Scale bar = 5 µm. **C. Histogram** summarizes the number of *Pomc*+ cells that co-express *Prl* or *Gh* mRNAs in adult pituitary. **Left graph:** histogram summarizes the number of dual positive cells (*POMC*^+^*Prl*^+^; 74%) and cells positive for *Pomc* but not *Prl* mRNA (*POMC*^+^/*Prl*^-^; 9.8%). **Right graph:** histogram summarizes the number of dual positive cells (*POMC*^+^*Gh*^+^; 59%) and cells positive for *Pomc* but not *Gh* mRNA (*POMC*^+^/*Prl*^-^; 18.3%).

### *Tshb* expression is primarily localized in the multi-hormone cluster rather than in a dedicated thyrotrope lineage

*Tshb* is generally considered to constitute one of the three Pou1f1-dependent pituitary hormone genes (Fig. S1) and the expression of *Tshb* is thought to define a distinct POU1F1-dependent ‘thyrotrope’ lineage (Zhu et al., 2007). While the scRNA-seq analysis defined clusters corresponding to the two major POU1F1-dependent lineages, somatotrope and lactotrope, as well clusters corresponding to all three of the POU1F1-independent lineages (corticotrope, melanotrope, and gonadotrope), the analysis failed to identify a discrete cell cluster that specifically corresponded to thyrotropes. Instead, *Tshb*+ cells mapped to all three of the the major ‘*Pou1f1* lineage’ clusters with a substantial representation in the multi-hormone cluster (Fig. 1C). A 2-dimensional scatter plot of *Tshb*, *Gh*, and *Prl* mRNA distributions revealed that 86.4% of *Tshb^+^* cells co-expressed both *Gh* and/or *Prl* mRNAs (Fig. 4A). This observation was validated by combining IF detection of TSHβ protein with scRNA FISH detection of *Gh* mRNA in male pituitaries; 84.6% of the TSHβ positive cells by IF were also positive for *Gh* mRNA by scRNA FISH (Figs. 4B **and** C). The co-expression of TSHβ and GH proteins was confirmed by IF in cells from an 8-week WT CD1 male mouse; 61% % of TSHβ+ cells were also positive for GH protein (Figs. S4C **and** D). These orthogonal studies lead us conclude that *Tshb* expression, while demonstrating the expected localization to *Pou1f1*^+^ cells, does not define a distinct ‘thyrotrope’ cell cluster. Instead, we find that *Tshb* is expressed in conjunction with *Gh* and/or *Prl*, and maps predominantly to the multi-hormone cluster.

**Figure 4.**
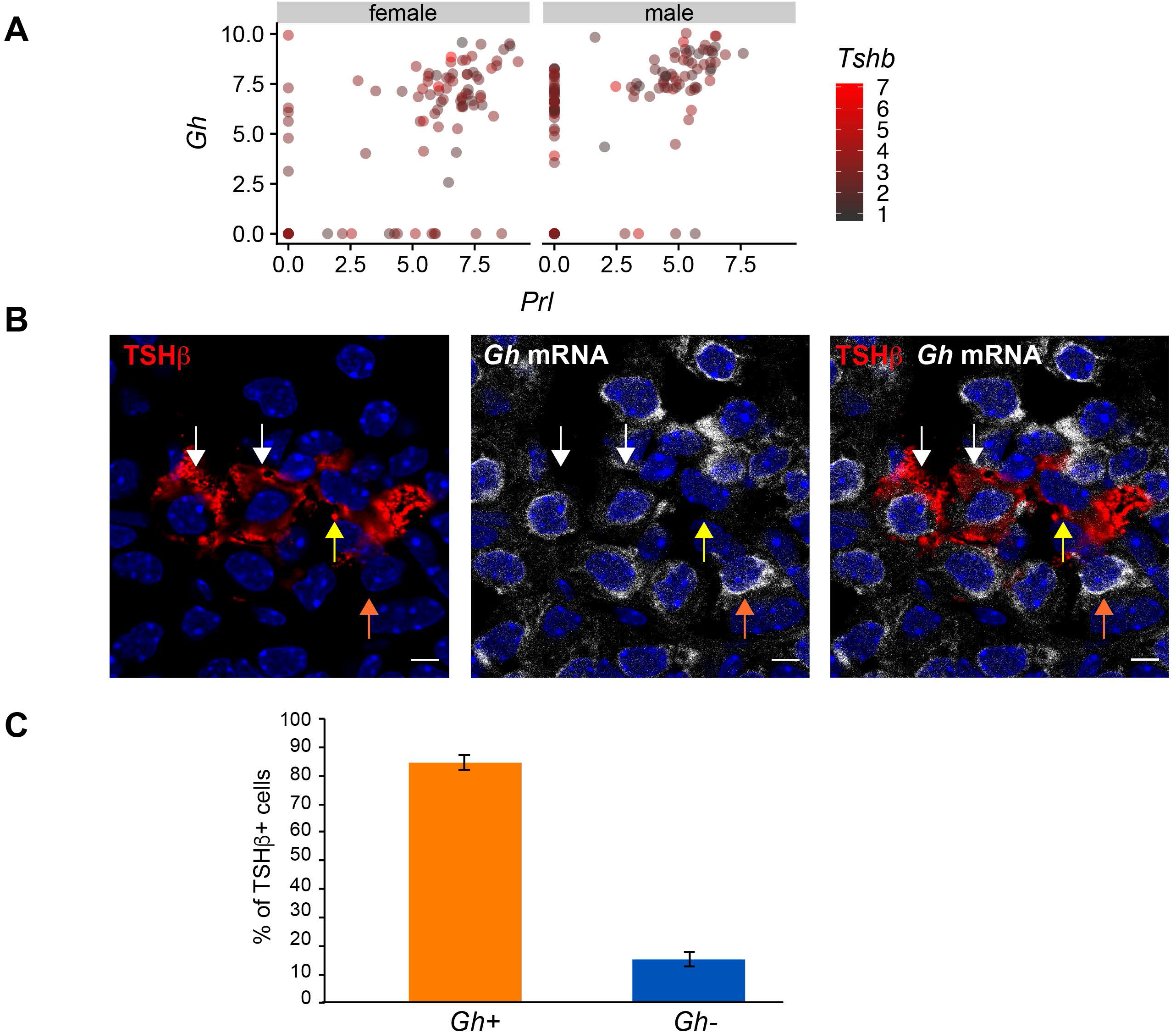
*Tshb* mRNA is predominantly co-expressed along with *Gh* and/or *Prl* mRNAs. **A. 2D scatter plot of *Gh* or *Prl* mRNAs in the *Tshb* mRNA expressing cells.** The analysis of Drop-seq data reveals that majority of the *Tshb*-expressing cells co-express *Gh* and/or *Prl* mRNAs. The levels of *Gh* and *Prl* mRNAs are represented on each of two labeled axes. The level of *Tshb* mRNA in each cell is indicated (heat map legend to the right). A small subset of cells express *Tshb* mRNA in the absence of *Gh* and/or *Prl* mRNAs [13.6% (30/220) in Drop-seq analysis]. **B. Prevalence of TSHβ and *Gh* co-expression as revealed by combined RNA-FISH/IF.** Detection of TSHβ protein (red in the left panel) was performed by IF (red in left panel) and *Gh* RNA detection was established by scRNA FISH (gray in the middle panel). The right panel is a merge of the RNA FISH and IF images. This single field analysis is representative of pituitary cells from tissue sections of three mice (two male mice and one female mouse). The white arrows indicate cells with robust expression of TSHβ protein and *Gh* mRNA. The orange arrow indicates a cell with robust expression of *Gh* mRNA but without TSHβ protein signal. The yellow arrow indicates a cell with TSHβ protein but without *Gh* mRNA signal. Nuclei were stained with DAPI (blue). Scale bar = 5 µm. **C. Summary of the analysis of cells expressing TSHβ and *Gh* mRNAs.** The RNA FISH/IF analysis was performed in three pituitaries (n = 3). The data reveals that 84.6% of the TSHβ protein positive cells co-expressed *Gh* mRNA.

### Sexual dimorphism in the organization and composition in the mouse pituitary

Synthesis and secretion of the pituitary hormones are impacted by multiple gender-specific developmental and physiologic stimuli. These controls drive critical somatic alterations linked to puberty, reproductive cycling, pregnancy, and lactation. The degree to which these gender specific functions are linked to differences in the cellular composition of the male and female pituitaries remains poorly defined (Lamberts and Macleod, 1990; Nishida et al., 2005). To address the issue of sexual dimorphism at the single-cell level we compared the transcriptomes of cells isolated from male and female pituitaries in our Drop-seq data set (Fig. 5A). Gender-specific analysis of *Gh* and *Prl* expression showed that *Gh*+/*Prl*-cells were more abundant in the male pituitary while the *Gh-*/Prl+ cells were more abundant in the female pituitary (Fig. 5B). Further analysis of major *Pou1f1*^+^ clusters revealed a relative predominance of somatotropes over lactotropes in males with a reciprocal enrichment of lactotropes over somatotropes in females (Fig. 5C). These data were consistent with the scRNA-FISH data (Fig. 2). The scRNA-FISH analysis of tissue sections confirmed this observation, revealing that the *Gh*-/*Prl*+ cells and *Gh*+/*Prl*-cells were relatively enriched in the females and males, respectively.

**Figure 5.**
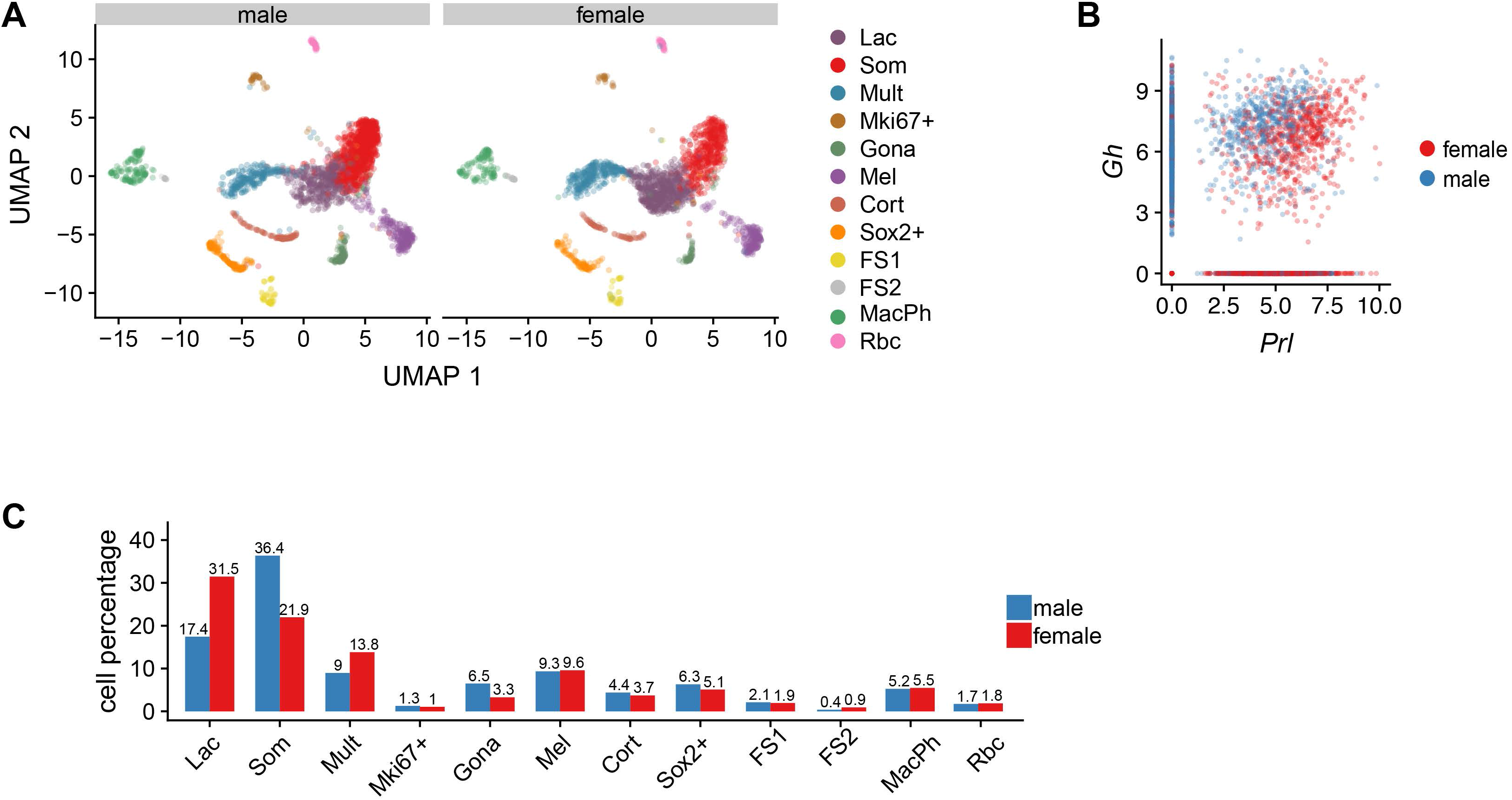
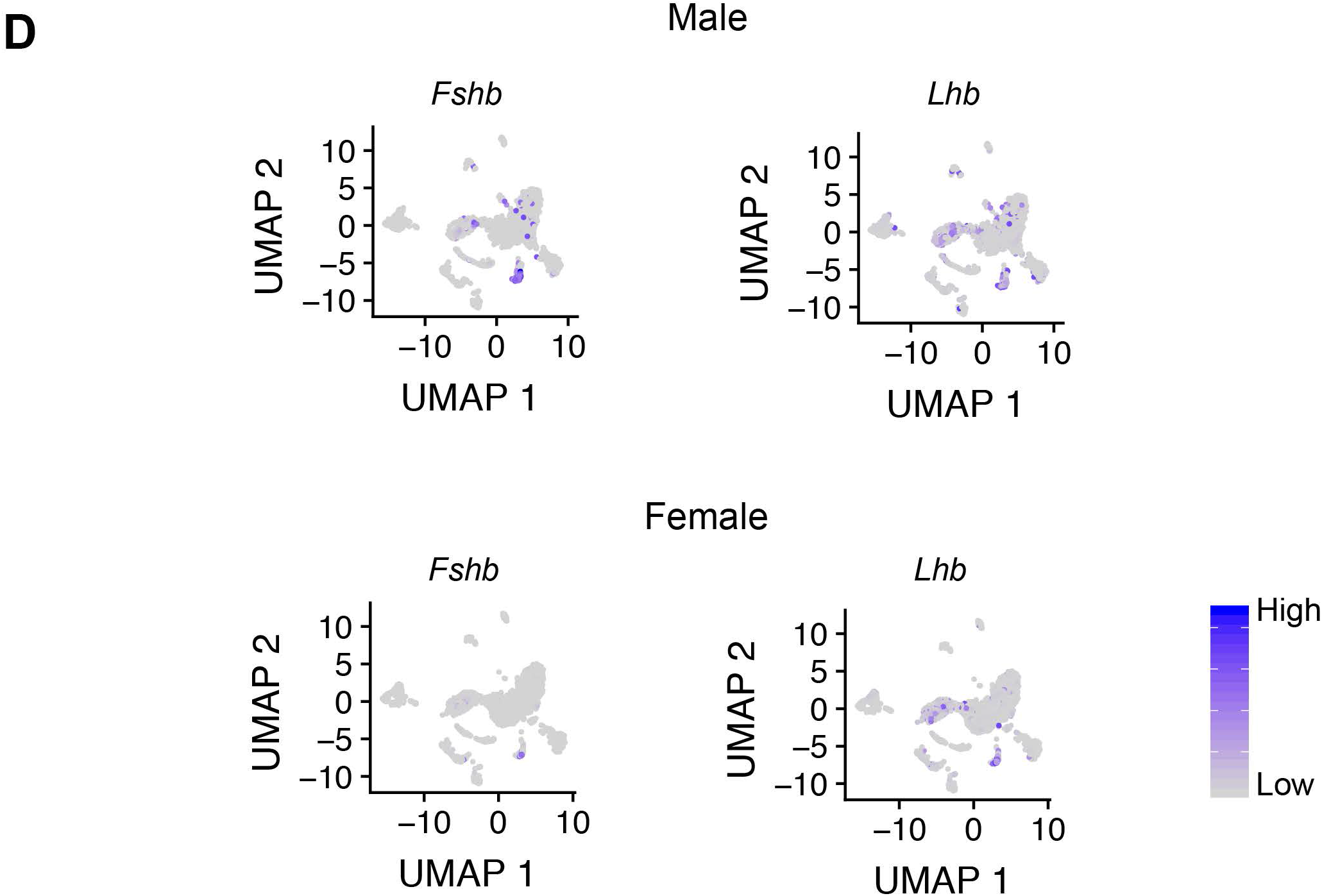
Clustering analysis of scRNA-seq data reveals sexual dimorphism in pituitary cell composition. **A. UMAP plot of cells isolated from age matched male (left panel) and female (right panel) pituitaries.** The clusters are color-coded as in Fig. 1. **B. 2D scatter plot of *Gh* or *Prl* mRNAs in the cells from the pituitaries of 8-week old WT mice.** The cells from different genders are labeled. The data reveal that the majority of the *Gh+* but *Prl-* cells are in the males while the majority of the *Prl+* but *Gh-* cells are in the females. The data is consistent with the data in figure 2. **C. Sexual dimorphism of the pituitary cell composition.** The histogram summarizes the relative distributions of cells in the male and female pituitary in each defined cluster, (derived from **A**.). The percent of total pituitary cell population represented in each cluster is shown above the respective bars. These data reveal relative enrichment for somatotropes in the male and a reciprocal enrichment for lactotropes in the female. Analysis of the *Pou1f1*-negative clusters reveals a significant enrichment of gonadotropes in the males. **D. Sexual dimorphism of gonadotrope hormones and the gonadotrope cluster.** The feature plots reveals differences in the expression patterns of the gonadotrope hormones (*Lhb* and *Fshb*) in male mice (top panels) vs female mice (bottom panels). The heat map legend is at the bottom right corner.

Marked sexual dimorphism was also evident in the gonadotrope cluster. This cluster constituted a substantially larger fraction of the pituitary cell population in male compared with female mice (6.5% vs 3.3% of total pituitary cells, respectively) (Fig. 5A **and** C). This dimorphism in gonadotrope cluster size in the male is consistent with previously-reported higher serum levels of FSHβ in male *vs* female mice (Michael et al., 1980). In contrast to the predominant localization of *Fshb* mRNA in the gonadotrope cluster, the expression of *Lhb* was equivalent between males and females and was predominantly expressed in the multi-hormone cluster in both sexes (Fig. 5D). These data lead us to conclude that relative representations of somatotropes and lactotropes populations differ between the two genders and that the two gonadotrope hormones, *Lhb* and *Fshb*, are differentially expressed in the male and female pituitaries both quantitatively and with relation to cell clustering.

### The response of hormone expressing clusters to major physiologic demands

We next sought validate and extend our analyses of pituitary cell populations to settings of defined physiologic stress. One of the most dramatic alterations in pituitary hormone production is the 10-15 fold increase in serum PRL levels during lactation in both mice and humans (Le Tissier et al., 2015). The molecular mechanisms underlying this increase are unclear (Castrique et al., 2010). In particular, it remains unclear to what extent this increase reflects an expansion of the lactotrope lineage and/or to alterations in hormone gene expression *per* lactotrope. To address this issue, we compared in two independent experiments the pituitary cell-type composition in three 13 week-old lactating female mice with an age-matched virgin female control ([Rep 1 (one mouse) and Rep 2 (two mice)] Fig. 6). This analysis revealed several noteworthy observations. First, the overall pattern of cell clustering in these 13 week old females was similar to that of 7-8 week old mice (male and female) with the formation of a multi-hormone cluster in both studies (compare Figs. 6A and 1A). Second, the comparison revealed an expansion of the lactotrope cluster in the lactating state (from 29% in control to 35.5% in lactating mice) with reciprocal decreases in representation of clusters representing multi-hormone, gonadotropes, corticotropes, and Sox2+ cells while the representation of somatotropes remained unchanged (Fig. 6B). These data demonstrate substantial shifts in specific pituitary cell populations in the transition to the lactating state with a net increase in the representation of lactotropes.

**Figure 6.**
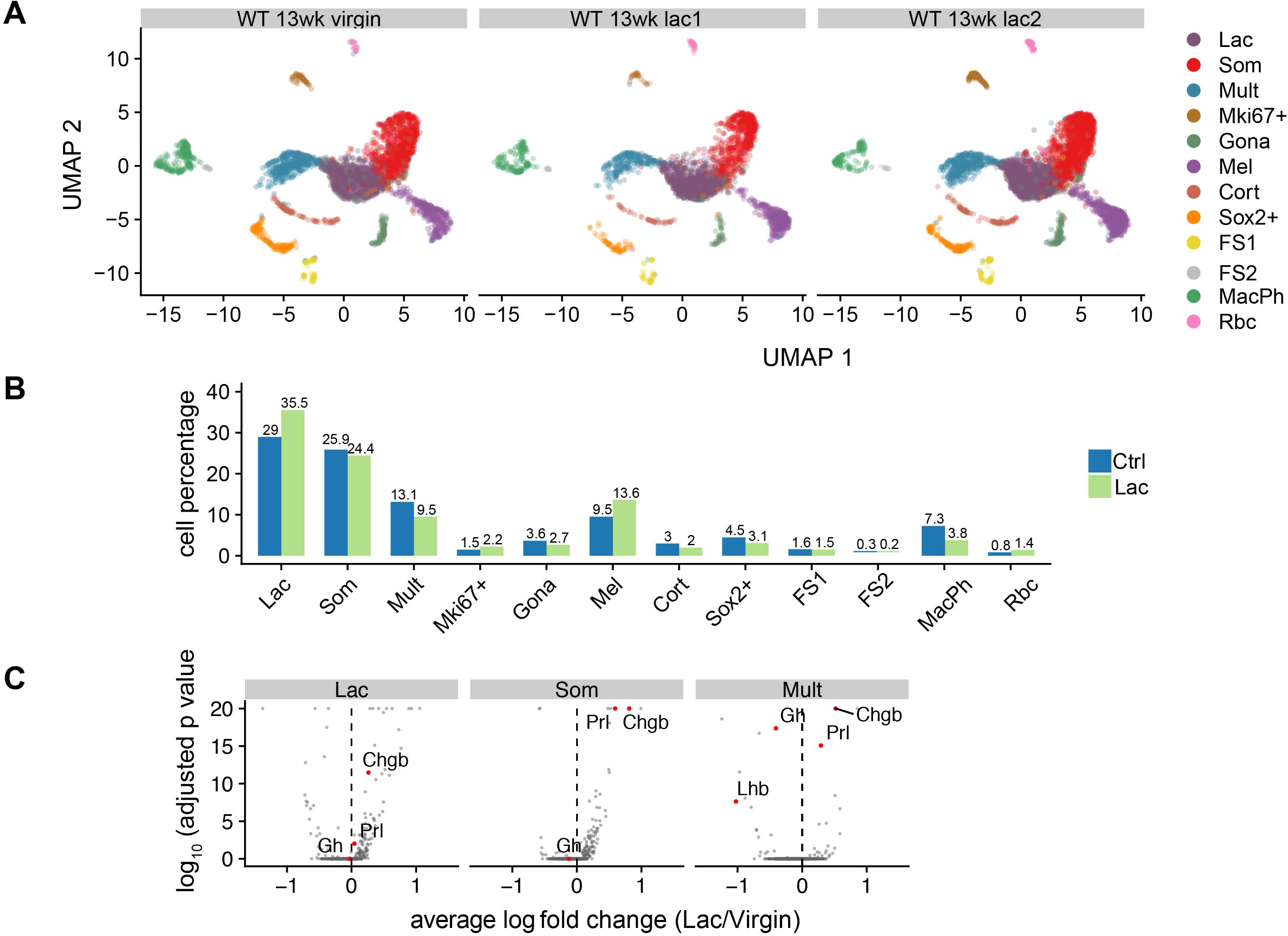
Single cell transcriptome analysis of the pituitary during lactation. **A. Clustering of adult pituitary cells based on principal component analysis of single cell transcriptomes.** The UMAP plot reflects the clustering analysis of individual adult pituitary cell transcriptomes (n = 11,566). The data represent the summation of pituitaries isolated from 13-week old female mice. Each color-coded cluster was assigned as in Fig. 1. **B. The pituitary cell composition in lactating females.** The histogram compares the cell representation in each indicated cluster between the age matched (13w) control vs the lactating (Lac) mice. The percent of total pituitary cell population represented in each cluster is shown above the respective bar. **C. Differential gene expression analysis of the *Pou1f1*^+^ lineages from lactating and sexual naïve control mice.** **Left panel: Analysis of lactotrope cluster.** The analysis was performed on 1,035 lactotropes from control and 2,750 lactotropes from lactating mice. The volcano plot shows the Log fold change (lactation/Virgin) in this value between the two groups on the X-axis and the significance (Log_10_ p-value) of the binary comparison for each gene on the Y-axis. Genes of particular interest (the hormones *Prl*, *Gh*, *Lhb*, and the neuroendocrine vesicle secretory protein, *Chgb*) are indicated as labeled red dots (details in text). **Middle panel. Analysis if somatotrope cluster.** The analysis was performed on 1,892 somatotropes from lactating mice compared with 924 somatotropes from an age matched female control. The volcano plot is organized as in left panel and highlights genes of particular interest (*Prl, Gh, and Chgb)* (details in text). **Right panel. Analysis of multi-hormone cluster.** The analysis compared the levels of expression of genes in the multi-hormone cluster in lactating females (n = 736) with an age matched female control (n = 469). Each dot represents mRNA expression from a specific gene. The volcano plot is organized as in left panel and highlights genes of particular interest (*Prl, Gh, Lhb,* and *Chgb*) (details in text)

The question of how lactation impacts *Prl* production was further explored by a focused analysis on the lactotrope transcriptomes. We performed PCA analysis on 3,785 cells in the lactotrope cluster (2,750 from lactating mice and 1,035 cells from age-matched control mice). The level of *Prl* mRNA was significantly higher in cells within the lactotrope cohort of cells originating from the lactating mice (adjusted *p*-value =0.0095) (Fig. 6C). This higher level of *Prl* mRNA levels in the lactotropes of the lactating mice was paralleled by an increase in expression of the neuroendocrine vesicle secretory protein *Chgb* (Barbosa et al., 1991; Gill et al., 1991) (adjusted *p*-value =3.36×10^-37^) (Fig. 6C). The increase in *Chgb* is consistent with the high demand for PRL secretion during lactation. These data lead us to conclude that the enhancement of PRL production during lactation reflects the combined effects of an increase in the lactotrope population as well as an elevation in *Prl* gene expression within the lactotrope cells.

In addition to its expression in lactotropes, *Prl* expression is also a prominent attribute of the multi-hormone cell cluster (Figs. 1 **and** 6C). For this reason, we next sought to determine whether the cells in the multi-hormone cluster contributed to the increased pituitary production of PRL in support of lactation. We noted that the representation of the multi-hormone cluster in the lactating mice decreased from that in the control (13.1% vs 9.5%, respectively) (Fig. 6B) while the expression of *Prl* in the cells within this cluster increased substantially (adjusted *p*-value = 8.42×10^-16^) (Fig. 6C). In contrast, the expression of *Lhb* in the cells within this cluster significantly decreased (adjusted *p*-value = 2.34×10^-8^). This is consistent with prior reports that the LHβ secretion is decreased during pregnancy in parallel with the inhibition of estrous cycle (Smith and Fox, 1984). We also observed a highly significant increase of *Prl* expression (adjusted *p*-value = 9.11×10^-126^) in the somatotropes of lactating mice compared with the control (Fig. 6C). In summary, these results demonstrated that the cells in the multi-hormone and somatotrope clusters undergo transcriptomic shifts that support increased production of PRL from lactotropes in the lactating female. These findings lead us to conclude that the induction of *Prl* expression in lactating females is based on shifts in cell composition along with shifts in cluster transcriptome profiles.

We next assessed the impact of a second major stress on the pituitary; the overexpression of *Ghrh*. GHRH is the major hypothalamic regulator of somatotrope function (Mayo et al., 1988); it is delivered to the anterior pituitary *via* the hypothalamic/pituitary venous portal circuit and binds to somatotrope cell surface receptors (GHRHR) to stimulate somatotropes proliferation of as well as to stimulate expression and secretion of GH (Gaylinn, 2002; Mayo et al., 2000). A well-described *mt/hGhrh* transgenic model (Mayo et al., 1988) has been previously reported and shown to drive high levels of GH expression in the mouse with resultant gigantism (Ho et al., 2002; Mayo et al., 1988). We performed Drop-seq analysis of 2,706 pituitary cells isolated from two 8-week old virgin female mice carrying the *mt/hGhrh* transgene (Mayo et al., 1988) and age-matched virgin non-transgenic females. This analysis revealed the assembly of an array of clusters that paralleled those detected in 8-week old non-transgenic mice as well as the 13-week old virgin and lactating females (Fig. 7A c/w Figs. 1 and 6). A multi-hormonal cluster was again observed in this analysis as well as the full array of hormone-expressing and non-hormonal cell clusters that were seen in the preceding analyses. When compared with the WT control, the analysis of the *hGhrh* transgenic pituitaries revealed a marked expansion in the somatotrope cluster; from 21.9% of total pituitary cells in WT to 33.8% in the *hGhrh* transgenic (Fig. 7B). This increase in the representation of the somatotrope cluster was paralleled by a reciprocal decrease in the representations of the multi-hormone (13.8% to 7.7%) and lactotrope (31.5% to 25.9%) clusters (Fig. 7B). Differential gene expression analysis of lactotropes in the *hGhrh*-transgenic pituitaries revealed significant increases in *Gh* mRNA (Fig. 7C). Importantly, *Gh* expression was significantly increased in the multi-hormone cells with reciprocal decreases in *Prl*, *Lhb*, and *Pomc* mRNAs were (Fig. 7C). These changes in cell cluster representation and gene expression in response to *Ghrh* overexpression highlight the contributions and coordination of the three major *Pou1f1*-expressing cell clusters in both their relative cell representations and in their gene expression profiles. These studies further validated the utility of scRNA-seq analysis in identifying shifts in cell representation and transcriptome expression profiles in response to physiologic demands.

**Figure 7.**
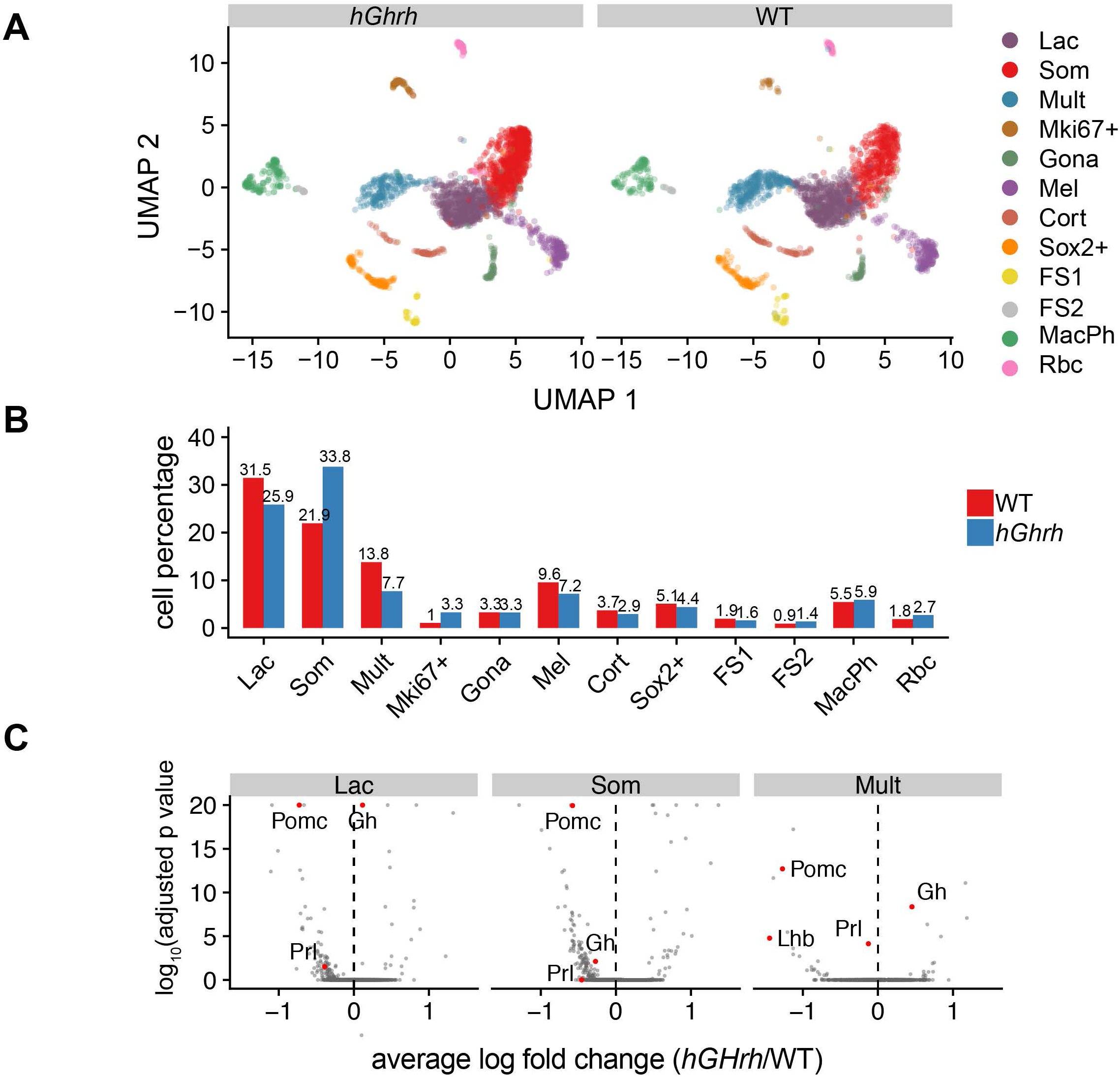
Single cell transcriptome analysis of the pituitary cells of mice overexpressing an *hGhrh* transgene. **A. Clustering of adult pituitary cells based on principal component analysis of single cell transcriptomes in *hGhrh* transgenic mice.** The UMAP plot reflects the clustering analysis of individual adult pituitary cell transcriptomes from four 8-week old female CD1 mice (Fig. 5A **left panel**) (n=2,110) and two 8-week old female *hGhrh* transgenic mice (n =2,706). Each color-coded cluster was assigned an ‘identity’ as in Fig. 1. **B. The pituitary cell composition in WT and *hGhrh* transgenic mice.** The histogram shows the cell compositions in the transgenic and 8-week old female CD1 (non-transgenic control; WT) mouse pituitaries, respectively. The representation of each total pituitary cell population, shown as a percent of total, is shown above the respective bar. **C. Differential gene expression analysis of the *Pou1f1*^+^ lineages in *hGhrh* transgenic mice** **Left panel. The comparison of the transcriptomes of lactotropes from WT and *hGhrh* transgenic mice.** The volcano plot (configured as in Fig. 8) shows the genes in which the expression was significantly altered during lactation. The positions of *Prl, Gh*, and *Pomc* are labeled in red (details see text). **Middle panel. The comparison of the transcriptomes of somatotropes from WT and *hGhrh* transgenic mice.** The positions of *Gh*, *Prl*, and *Pomc* are labeled in red (details see text). **Right panel. The comparison of the transcriptomes of multi-hormone cluster from WT and *hGhrh* transgenic mice.** The positions of *Prl*, *Gh*, *Lhb* and *Pomc* are labeled in red (details see text).

## Discussion

Current understanding of pituitary function is based upon a binary model in which each of the six distinct endocrine cell types is dedicated to the synthesis and secretion of its corresponding polypeptide hormone (Fig. S1). The function of each lineage is considered as mutually exclusive and under the control of a corresponding set of discrete hypothalamic/pituitary regulatory circuits (Davis et al., 2013; Kelberman et al., 2009; Zhu et al., 2007). However, scattered reports of pituitary cells co-expressing multiple hormones have suggested that the organization and function of the pituitary may be more complex than commonly appreciated (Childs, 2000; Frawley and Boockfor, 1991; Seuntjens et al., 2002a; Villalobos et al., 2004b) and the prevalence and complexity of ‘multi-hormone’ cells in the adult anterior pituitary has remained undefined on a systematic and global level. Thus, understanding the compositions and functions of pituitary cell lineages and their relationships to hormone expression warrants further study.

Here we report a series of orthogonal single-cell analyses that address fundamental aspects of pituitary cell organization, lineage specification, and functional plasticity. Our single cell transcriptome analysis yielded findings that were consistent with certain aspects of existing models while substantially extending and modifying others. While the analysis confirmed the presence of three discretely clustering *Pou1f1*-independent cell lineages (corticotropes, melanotropes, and gonadotropes (Fig. 1A)), it reveals that the organization and functions of the Pou1f1-dependent lineages are substantially complex (Fig. 1). The initial presumption going into this study was that the clustering analysis would generate three *Pou1f1*^+^ clusters corresponding to somatotropes, lactotropes, and thyrotropes, each dedicated to the exclusive expression of its corresponding hormone; *Gh, Prl,* and *Tshb*, respectively (Fig. S1) (Zhu et al., 2007). However, our unbiased scRNA-seq failed to confirm the existence of a unique ‘thyrotrope’ lineage and instead revealed that the majority of *Tshb* mRNA expression, along with a significant fraction of *Gh* and *Prl* mRNAs, were derived from a novel *Pou1f1*-expressing ‘multi-hormonal’ cell cluster. The discovery of this multi-hormone cell cluster challenges current models of pituitary lineage distinctions and the segregation of hormone expression.

Our multiple independent scRNA-seq analyses on pituitaries isolated from mice of varying ages and genders as well as under two well defined setting of physiologic stress reproducibly identified the unique multi-hormone cluster (Figs, 1, 6**, and** 7). Most intriguing was the observation that the cells in this cluster not only expressed well defined POU1F1-dependent genes such as *Gh*, *Prl*, and *Tshb*, but also expressed robust levels of mRNAs encoded by two genes, *Pomc* and *Lhb*, that as traditionally categorized as POU1F1-independent. A series of single cell RNA and protein imaging studies documented co-expression of the ‘POU1F1-dependent’ and the ‘POU1F1-independent’ hormone genes within individual cells (Figs. 2, 3, S4, and S5).

While multi-hormone producing cells have been previously identified by others using immuno-fluorescent and targeted RT-PCR analyses (Seuntjens et al., 2002a; Seuntjens et al., 2002b; Villalobos et al., 2004a), their abundance, array of hormone gene expression, and physiological function(s) have remained unclear. Our analyses reveal that the cells in this multi-hormone cluster can be defined by a unique transcriptional signature and respond to physiological demands *in vivo*. For example, in lactating females, the level of *Prl* expression increases in the cells in this cluster with reciprocal decreases in *Lhb* and *Gh* mRNAs. Likewise, in transgenic mice overexpressing the *hGhrh* gene, *Gh* expression increases in the multi-hormone cells while the expression of other pituitary hormones significantly decrease. These observations suggest that cells in the multi-hormone cluster are able to respond to substantial physiological demands and exhibit plasticity of pituitary function and hormone production.

The multi-hormone cluster constitutes a substantial fraction (11.1%) of total pituitary cell population. The cells within this cluster appear to serve as a transitional cell pool that can respond to shifts in hormone production. Such shifts are likely essential to support physiological demands imposed by pregnancy, lactation, and pubertal growth. Changes in hormone gene expression patterns in such large number of cells can significantly facilitate the response of the hypothalamic/pituitary axis to acute physiologic demands. It will be of interest to further explore the molecular/signaling pathways that control the cellular plasticity of the cells within the multi-hormone cluster.

The developmental origin(s) of the *Pou1f1^+^* multi-hormonal cells in adult mouse pituitary is presently unclear. The broad functional capacity of these cells cannot be accounted for in current models of anterior pituitary development and cellular differentiation (Fig. S1). However, recent studies suggest that these cells may be induced by the actions of PROP1. PROP1, a paired-like homeodomain transcription factor, was initially described as a factor that triggers the activation of *Pou1f1* and the differentiation of the POU1F1-dependent lineages during pituitary development (Andersen et al., 1995; Gage et al., 1996). However, recent lineage tracing studies in the mouse reveal that PROP1 is in fact expressed in progenitor cells that generate the full spectrum of endocrine cell types in the pituitary, those considered to be ‘POU1F1-dependent’ as well as those considered to be ‘POU1F1-independent’ (Davis et al., 2016; Perez Millan et al., 2016). Multi-hormone expressing cells may therefore be generated through the activity of PROP1 during pituitary development.

In addition to revealing the presence of the multi-hormone cell cluster, the Drop-seq analysis highlighted complex relationships between the functions of the two other major POU1F1 clusters, the somatotropes and lactotropes. While the cells in these two clusters could be readily identified by a preponderant expression of either *Gh* or *Prl* as well as by a number of corresponding signatory mRNAs (see Fig, 1 and accompanying text), the majority of these cells co-expressed *Gh* and *Prl* to varying degrees (Fig. 1 and Fig. 2). Furthermore, the levels of *Prl* expression in somatotropes increased substantially in lactating mice while *Gh* expression increased significantly in the lactotropes of the *hGhrh* transgenic mice (Fig. 6 and 7). These data suggest that the capacity of the somatotrope and lactotrope lineages to express both *Prl* and *Gh* may be critical to a robust response to major physiological demands.

The synthesis and secretion of the pituitary hormones are impacted by multiple physiologic variables that directly relate to sexual maturation, reproduction, and somatic development. As such, sexual dimorphism in pituitary structure and hormonal output has been previously identified in a number of these settings (Lamberts and Macleod, 1990; Michael et al., 1980; Nishida et al., 2005). Of interest, the Drop-seq analysis of the pituitaries of 7-8 week old, sexually-naïve mice confirmed that even at the young adult stage there is strong evidence for sexual dimorphism. The most striking gender distinctions in this regard was the dominance of the somatotropes in the males (Fig. 5), the relative enrichment of lactotropes in in females (Fig. 5), and the distinct patterns of gonadotrope hormone expression between the two genders (Fig. 5).

In conclusion, we have combined single-cell RNA sequencing with RNA and protein single cell imaging to analyze the transcriptomes of cells within the adult mouse pituitary. The results have revealed unanticipated levels of cellular diversity and lineage plasticity in pituitary cell-type composition and hormone expression. The findings reveal a significant fraction of cells that co-express multiple hormones, sexual dimorphisms of lineage composition and cell prevalence, and the plasticity of cell functions in response to major physiologic demands. These single-cell transcriptomic data sets, along with experimental approaches to identifying the factors that underlie these complexities of lineage structure and function, can now be further extended to explore pituitary functions in settings of physiologic stress and disease.

## Methods

### Animals

Six-week old CD1 mice were purchased from Charles River (Wilmington, MA) and were housed for a minimum of 1 to 2 weeks in rooms with 12-hour light/dark cycles prior to studies. All aspects of the mouse studies were approved by the University of Pennsylvania Laboratory Animal Use and Care Committee. The *mt/hGhrh* transgenic mice were originally obtained from Dr. Ralph Brinster at the University of Pennsylvania (Hammer et al., 1985) and have been described in a number of our prior reports (Ho et al., 2002; Ho et al., 2008).

### Single cell preparation for Drop-seq

Single cell pituitary suspensions were prepared by non-enzymatic methods as previously descripted (Ho et al., 2011) with minor modifications to adapt to the Drop-seq protocol (Macosko et al., 2015). Briefly, the pituitaries were isolated and washed with cold PBS, incubated with 1 ml of enzyme-free cell dissociation buffer (Life technologies, Carlsbad, CA) for 1 minute, passed through a 40 µm cell strainer, and then re-suspend in 10 ml of PBS. The cells were diluted to 100 cells/µl in PBS with 0.01% BSA prior to the capture of the cells

### Single-cell RNA-Seq library preparation and sequencing

Drop-seq was performed as previously described with minor modifications (Macosko et al., 2015). Specifically, cells were captured on barcoded beads, reverse transcribed, and treated with exonuclease prior to amplification. The cDNA from an aliquot of 6,000 beads was amplified by PCR in a volume of 50 µL (25 µL of 2x KAPA HiFi hotstart readymix (KAPA biosystems), 0.4 µL of 100 µM TSO-PCR primer (AAGCAGTGGTATCAACGCAGAGT), 24.6 µL of nuclease-free water) to determine an optimal number of PCR cycles for cDNA amplification. The thermal cycling parameter was set to: 95°C for 3 min; 4 cycles of 98°C for 20 sec, 65°C for 45 sec, 72°C for 3 min; 9 cycles of 98°C for 20 sec, 67°C for 45 sec, 72°C for 3 min; 72°C for 5 min, hold at 4°C. The amplified cDNA was purified twice with 0.6x SPRISelect beads (Beckman Coulter) and eluted with 10 µL of water. 10% of amplified cDNA was used to perform real-time PCR analysis (1 µL of purified cDNA, 0.2 µL of 25 µM TSO-PCR primer, 5 µL of 2x KAPA FAST qPCR readymix, and 3.8 µL of water) to determine the additional number of PCR cycles needed for optimal cDNA amplification (Applied Biosystems QuantStudio 7 Flex). PCR reactions were then optimized per total number of barcoded beads collected for each Drop-seq run, adding 6,000 beads per PCR tube, and run according to the aforementioned program to enrich the cDNA for 4 + 12 to 13 cycles. The amplified cDNA was ‘tagmented’ using the Nextera XT DNA sample preparation kit (Illumina, cat# FC-131-1096), starting with 550 pg of cDNA pooled in equal amounts from all PCR reactions for a given run. After quality control analysis using a Bioanalyzer (Agilent), libraries were sequenced on an Illumina NextSeq 500 instrument using the 75-cycle High Output v2 Kit (Illumina). The library was loaded at 2.0 pM and the Custom Read1 Primer (GCCTGTCCGCGGAAGCAGTGGTATCAACGCAGAGTAC) was added at 0.3 µM in position 7 of the reagent cartridge. The sequencing configuration was 20 bp (Read1), 8 bp (Index1), and 50 or 60 bp (Read2). Two male samples and two female samples (two mice per sample) from 8-week old CD1 mice, 1 sample from 13-week old virgin mouse, 2 samples from 13-week lactation mice and 1 sample from two *mt/hGRF* transgenic mice (**Table S2**), were analyzed with Drop-seq in five sequencing runs.

### Read mapping

Paired-end sequencing reads of Drop-seq were processed as previous described (Hu et al.). Briefly, after mapping the reads to the mouse genome (mm10, Gencode release vM13), exonic reads mapped to the predicted strands of annotated genes were retrieved for the cell type classification. Uniquely mapped reads were grouped by cell barcode. To digitally count gene transcripts, a list of UMIs in each gene, within each cell, was assembled, and UMIs within ED = 1 were merged together. The total number of unique UMI sequences was counted, and this number was reported as the number of transcripts of that gene for a given cell. Raw digital expression matrices were generated for all of the 8 samples.

### Cell type classification

To enable directly comparative analyses among different conditions, such as male versus female, lactation versus virgin, WT versus *mt/hGRF* transgenic mice, we used Seurat 3 (v. 3.0.0) which has been demonstrated as an effective approach to perform joint analyses and build an integrated reference (Stuart et al., 2019b) (Figs. S2A **and** S2B). The raw digital expression matrices of all 8 samples from Drop-seq runs were combined and loaded into the Seurat 3. For normalization, UMI counts for all cells were scaled by library size (total UMI counts), multiplied by 10,000, and transformed to log space. Only genes found to be expressing in >10 cells were retained. Cell with a high percentage of UMIs mapping to mitochondrial genes (>=0.1) were discarded. In addition, cells with fewer than 300 UMI counts, fewer than 100 detected genes, or more than 4,000 detected genes were discarded, resulting in 19,867 cells from 8 samples. The nUMIs and nGenes are shown (Fig. S2C). The top 2,000 highly variable genes (HVGs) were identified using the function *FindVariableFeatures* with “vst” method. Canonical correlation analysis (CCA) was used to identify common sources of variation among WT, *hGHRF* transgenic mice, and lactating mice. The first 30 dimensions of the CCA was chosen to integrate the 6 datasets, including 2 replicates of 8-week old WT mice, 1 replicate of 13-week old virgin female mice, 2 replicates of 13-week old lactation female mice and 1 replicate of 8-week old *mt/hGRF* transgenic mice. After integration, the expression levels of HVGs in the cells were scaled and centered along each gene and was subjected to PCA analysis. The top 25 PCs were selected and used for 2-dimension reduction by Uniform Manifold Approximation and Projection (UMAP). Clusters were identified using the function *FindCluster* in Seurat with the resolution parameter set to 0.4. Assessing a number of different PCs for clustering revealed that the variation of PC number selection was relatively insensitive to the clustering results (Fig. S2D). Cells were classified into 14-22 clusters with the resolution parameter from 0.4 to 1 (Fig. S2E). Clustering resolution parameters were varied quantitatively based on the number of cells being clustered. After the clustering results with different resolutions were compared and evaluated, we chose a resolution value of 0.4. Using this approacg we were able to assign 19,660 cells to 14 clusters. We further filtered out two clusters; one cluster contained 638 cells with low quality and the other cluster, with almost double number of genes per cell as compared to other cell clusters, were considered cell doublets. In all, 1,070 cells (5.3% of input data) were removed from the downstream analysis and 18,797 cells were assigned into 12 cell clusters. Marker genes were identified using the function *FindMarkers* in Seurat. Cell type was annotated based on top ranked marker genes. To this end, the information was used to build an integrated cell type reference of mouse pituitary (**Table S2**).

The single cell suspension of 8-week old mouse pituitary was loaded onto a well on a 10x Chromium Single Cell instrument (10x Genomics). Library preparation was performed according to the manufacturer’s instructions. For cell type classification of 10xgenomic data set, low quality cells were filtered out using the same criterion as Drop-seq analysis pipeline (100 <nGenes < 4000 and nUMI > 300). We projected the PCA structure of Drop-seq built reference onto the 10x genomic dataset using *FindTransferAnchors* with the parameter “dims=1:25” in Seurat. *TransferData* function was used to classify the cell types of 10x genomic data set based on the reference data. 3,929 cells in 10xgenomic data set were assigned with cell type information (**Table S2 and** Fig. S3).

### Background correction of the cell transcriptomes

After clustering, we found that highly transcribed hormone genes (*Gh*, *Prl* and *Pomc*) could be detected in blood cells (Fig. S2F). This observation most likely represented cross-contamination of free RNAs. Since these mRNAs are highly expressed in the pituitary but should be absent in the blood cells, their levels in blood cells represent a background noise signals. We adapted a previous background correction approach (Han et al., 2018) to correct the level of contaminated mRNA. We assumed that the cell barcodes with less than 100 UMI or 100 nGene correspond to the empty beads exposed to free RNA during the cell dissociation or droplet breakage steps. The average of nUMI for the cell barcodes (< 100 Gene or < 100 UMI) was therefore calculated and used to perform a background correction by subtracting this value from the raw nUMI digital expression matrix, and the negative value was then set to zero. After correction, UMI counts for all cells were scaled by total UMI counts, multiplied by 10,000, and transformed to log space. We used the background corrected date set to generate feature plots and violin plots (Figs. 1B, 1C, 4A, 5B**, and** 5D).

### Identification of Differentially Expressed Genes

Differential gene expression analysis between control mice and *Ghrh* transgenic mice or between control mice and lactating mice was performed using the function *FindMarkers* in Seurat, using a Wilcoxon rank sum test. Genes with an adjust *P*-value less than 0.05 were considered to be differentially expressed.

### Probe design for single-cell RNA fluorescent *in situ* hybridization (scRNA FISH)

The strategy for the scRNA FISH probe design followed the strategy generally used for Single-molecule Oligopaint FISH studies (Beliveau et al., 2015). Each 40 nt primary probe contained 20 nt complementary to the exons of the mRNAs (*Gh*, *Prl*, and *POMC)* with a 20 nt non-genomic tag located at the 5’ end. 20 distinct primary probes scanning each targeted mRNA were used in each probe library. The tag sequence was unique for each mRNA library (Beliveau et al., 2015). A fluorophore-labeled secondary oligo with base complementarity to the tag sequence was used to detect the hybridized primary probes. All oligos were synthesized by Integrated DNA Technologies (IDT; Coralville, IA). The sequences of the primary and secondary oligos are available upon request.

### Preparation of pituitary tissue sections

The pituitaries of 8-week old mice were excised and washed in ice-cold PBS. The tissue was fixed in 4% paraformaldehyde for 2 hours at 4°C. The tissue was cryo-protected in 30% sucrose and embedded in Tissue-Tek OCT compound (Sakyra Finetek USA Inc, REF:4583) and frozen at −80°C. Coronal or sagittal sections (6 µm thickness) were generated and mounted on slides. The slides were dried at room temperature for 2 hours and stored at −20°C prior to analysis.

### scRNA FISH and immuno-fluorescent-scRNA (IF-scRNA) FISH

The procedures for RNA FISH were as described (Beliveau et al., 2015; Ho et al., 2013; Ragoczy et al., 2006) with modifications. Briefly, slides were removed from −20°C storage and dried at room temperature for 1 hour. The slides were washed with PBS, fixed with 3.7% formaldehyde in PBS for 20 minutes, washed with PBS and treated with 70% ethanol at 4°C for overnight. 500 pmol of primary oligo library (19 to 22 oligos) and equal amount of fluorophore-labeled secondary were used for each slide (6-8 tissue sections). Prior to hybridization, the slides were washed with PBS, sequentially dehydrated in 70%, 90%, and 100% ETOH, and then equilibrated in 10% formamide/2X SSC, pH 7.0. The mixture of the primary oligo probes and the secondary probes were hybridized to the cells in 10% formamide/10% dextran sulfate/2XSSC/5 mM ribonucleotide vanadate complex/0.05% BSA/1μg/μl *E.coli* tRNA. The probes were heat denatured at 85°C for 5 min., pre-annealed at 37°C, and then hybridized overnight at 37°C in a humidified chamber. Slides were sequentially washed in 10% formamide/2X SSC, pH 7 followed by 2X SSC at 37°C and then mounted with Fluoroshield with DAPI (Sigma, St. Louis, MO).

For IF analysis, the slides were removed from −20°C, dried at room temperature for 1 hour, washed with PBS containing 0.1% Triton X-100 (PBST) (3x 15 minutes), incubated with blocking buffer (2% BSA, 0.1% Triton X-100, and 5% normal donkey serum in PBS) for 1 hour and then incubated with primary antibodies for overnight at 4°C. GH was detected using monkey anti-rat GH that cross-reacts with mGh. PRL was detected using rabbit anti-mouse PRL (National Hormone and Peptide Program, NIH) or goat anti-PRL antibody (ThermoFisher Scientific, PA5-47140). ACTH and TSHβ were detected using rabbit-anti-ACTH antibody or TSHβ antibody, respectively (National Hormone and Peptide Program, NIH). The secondary antibodies used were donkey anti-human, anti-rabbit, anti-goat antibodies (Jackson immnoResearch Inc). The slides were mounted as for the scRNA FISH.

For the combined IF/scRNA FISH studies, the RNA FISH was performed as described above and the slides were then equilibrated in PBST for 10 minutes at room temperature, blocked with blocking buffer containing 2 mM ribonucleotide vanadate complex, and IF was performed as described above.

For the scRNA FISH and IF studies of disassociated single cells, the pituitaries were disassociated as described in the Drop-seq procedure followed by fixation in poly-lysine coated slides and the procedures for RNA and protein analyses were then performed as described above.

### Image Analysis

Image capture were collected on an Leica TCS SP8 confocal microscope platform. The images were analyzed using ImageJ software (NIH).

## Acknowledgments

We thank Dr. Sonny Nguyen for technical suggestions on scRNA FISH. We are grateful to Dr. Andrea Stout of the Microscopy Core in the Department of Cell and Development Biology, Perelman School of Medicine, University of Pennsylvania for help with the image capture. This work was supported by National Institutes of Health (NIH) Grants DK107453 (to S.L and Y.H) and R00HG007982, DP2HL142044 and a University of Pennsylvania Epigenetics Institute Pilot Grant (to H.W).

## Author Contributions

Y.H., H.W., and S.A.L designed experiments. Y.H., P.H., M.P, and S.C conducted the experiments. P.H., H.W., and P.G-C performed the bioinformatic analyses. Y.H. S.C and D.E. designed and performed the RNA-FISH in tissue sections. Y.H., P.H., H.W., and S.A.L wrote the manuscript. All authors read and approved the manuscript.

## Supplemental Figure Legends

**Figure S1.**
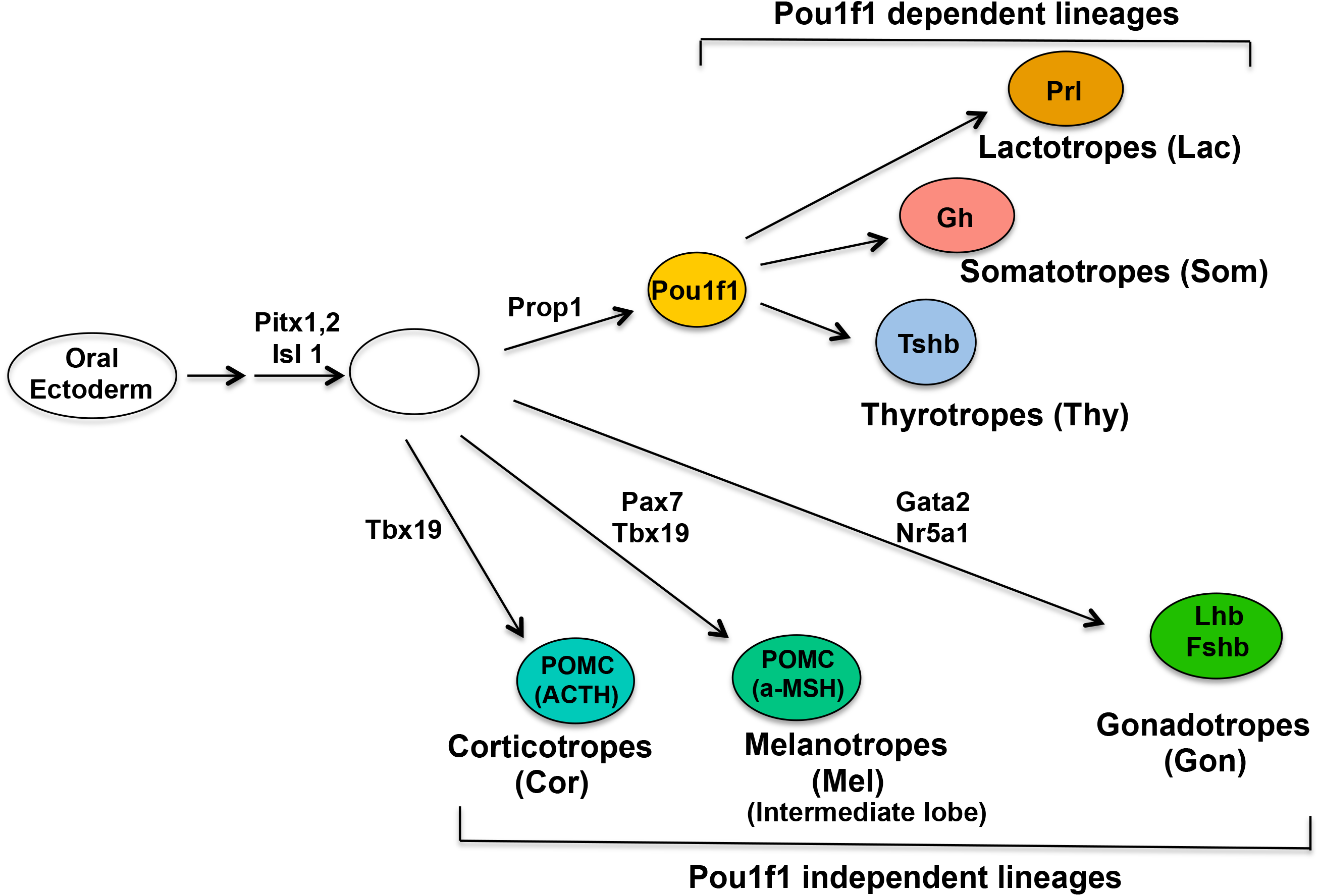
Standard model of anterior pituitary development and lineage specification. This diagram summarizes the standard model of pituitary lineage differentiation. The development of the mouse anterior pituitary from the oral ectoderm is driven by a complex array of signaling pathways and transcription factors, a subset of which are indicated. The adult pituitary is characterized by the presence of six terminally differentiated cell types, each defined by the expression of its corresponding hormone (Prl, prolactin; Gh, growth hormone; Tshb; β subunit of thyroid stimulating hormone; Lhβ and Fshβ, the β subunits of the luteinizing and follicle stimulating hormones; POMC, proopiomelanocortin prohormone; ACTH, adrenocorticotropic hormone; α-MSH, melanocyte stimulating hormone). Differentiation of three lineages (lactotropes, somatotropes and thyrotropes) is specifically driven by, and dependent on, the POU-homeo domain transcription factor, Pou1f1. The remaining three lineages (corticotropes, melanotropes, and gonadotropes) are considered to be Pou1f1-independent.

**Figure S2.**
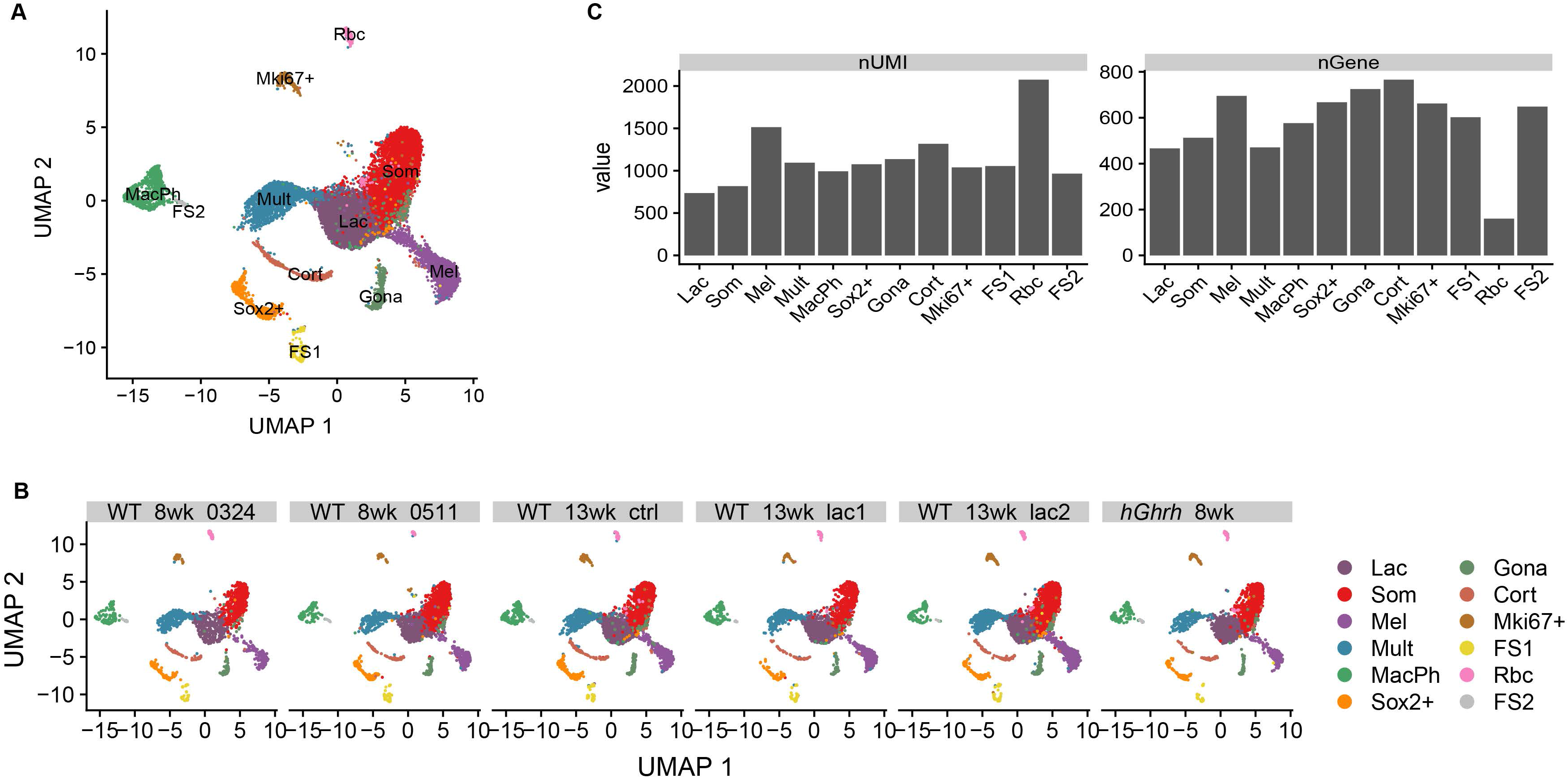
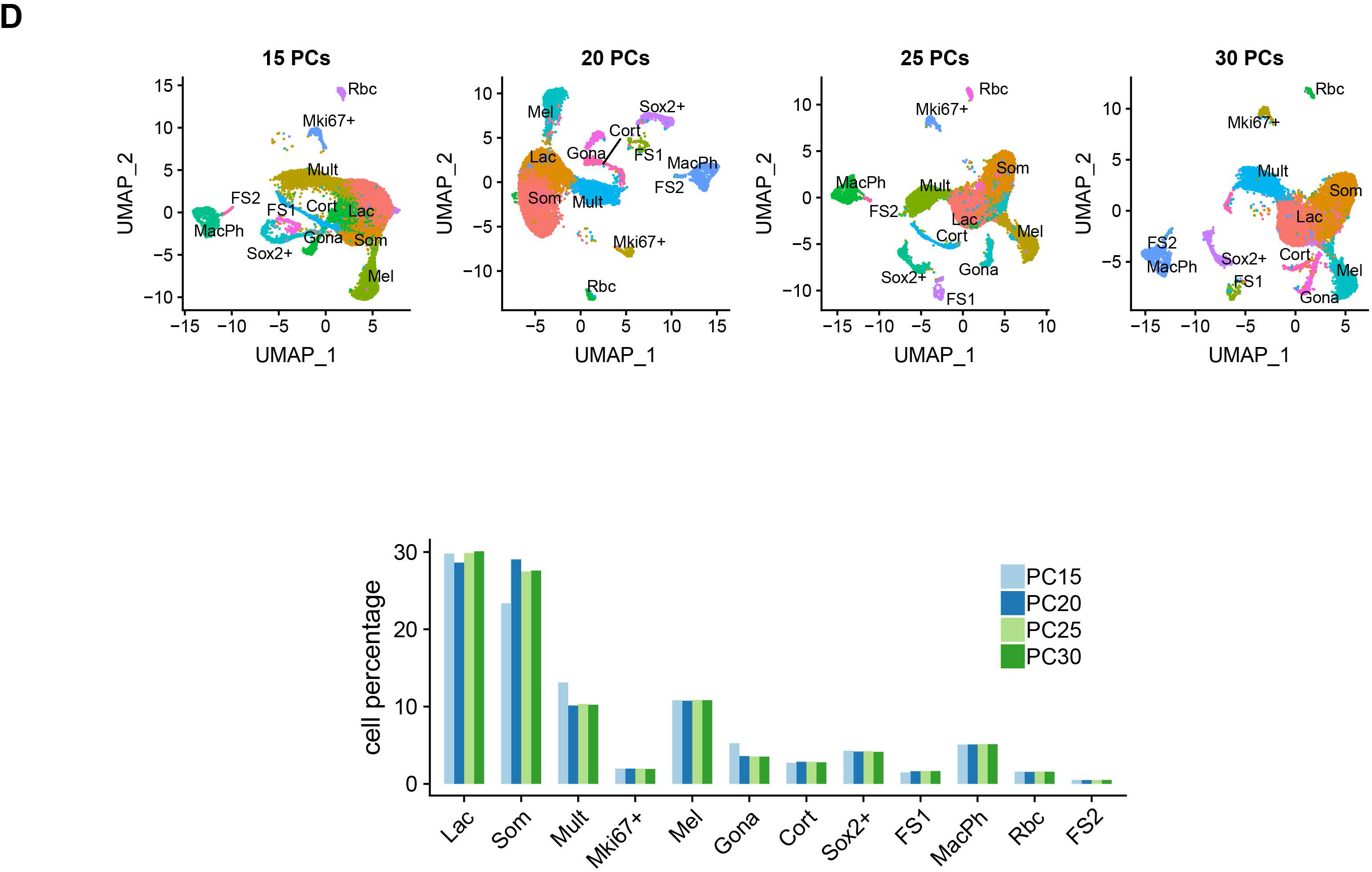
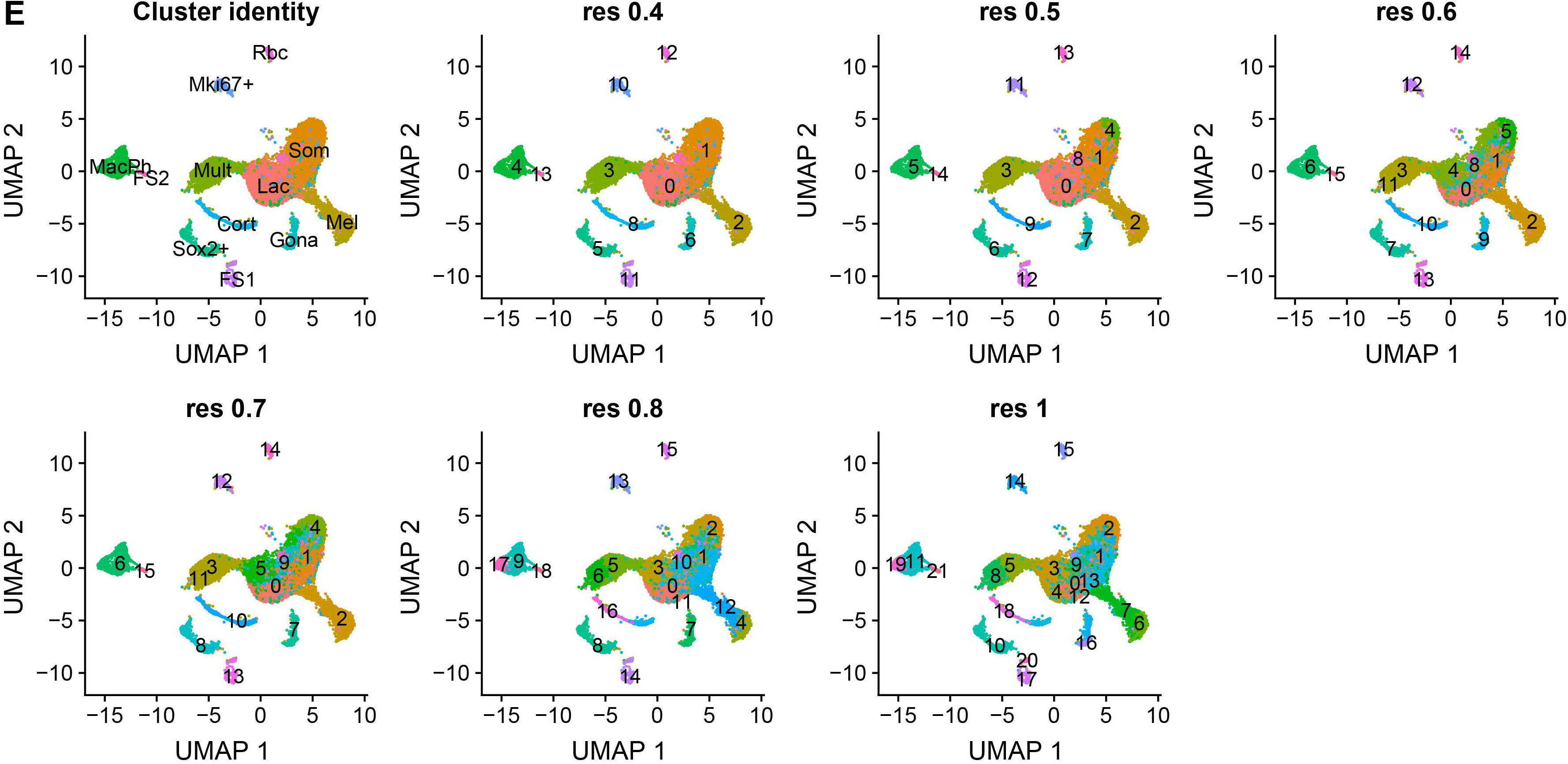
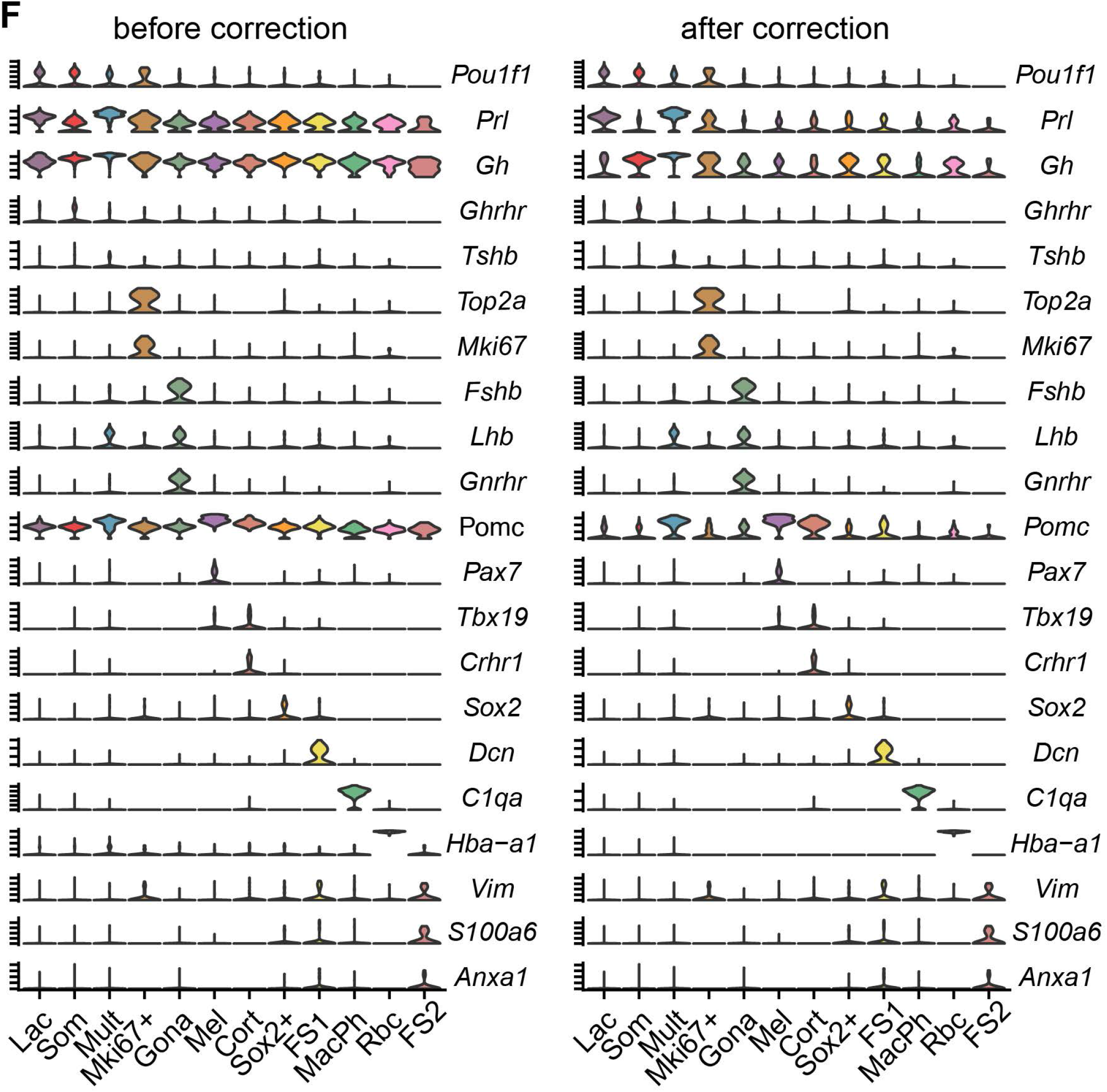
Cell type classification and background correction. **A. UMAP projections of 18,797 cells generated by integrating data sets of Drop-seq runs.** **B. UMAP projections across different replicates, ages, and physical conditions.** The clusters are color-coded and the identities of the clusters are labeled to the right of the UMAP plots. **C. Bar plot shows the average nGene and nUMI across cell clusters.** **D. Robustness of cell type classification to variation of number of PCs.** UMAP plots using varying PC numbers as shown above each respective the plot. Bar plot shows the effect of varying PC numbers on the cell type composition. **E. UMAP plots show the effect of varying resolution parameters on the number of cell clusters.** **F. Violin plots show the expression of marker genes before and after background correction.**

**Figure S3.**
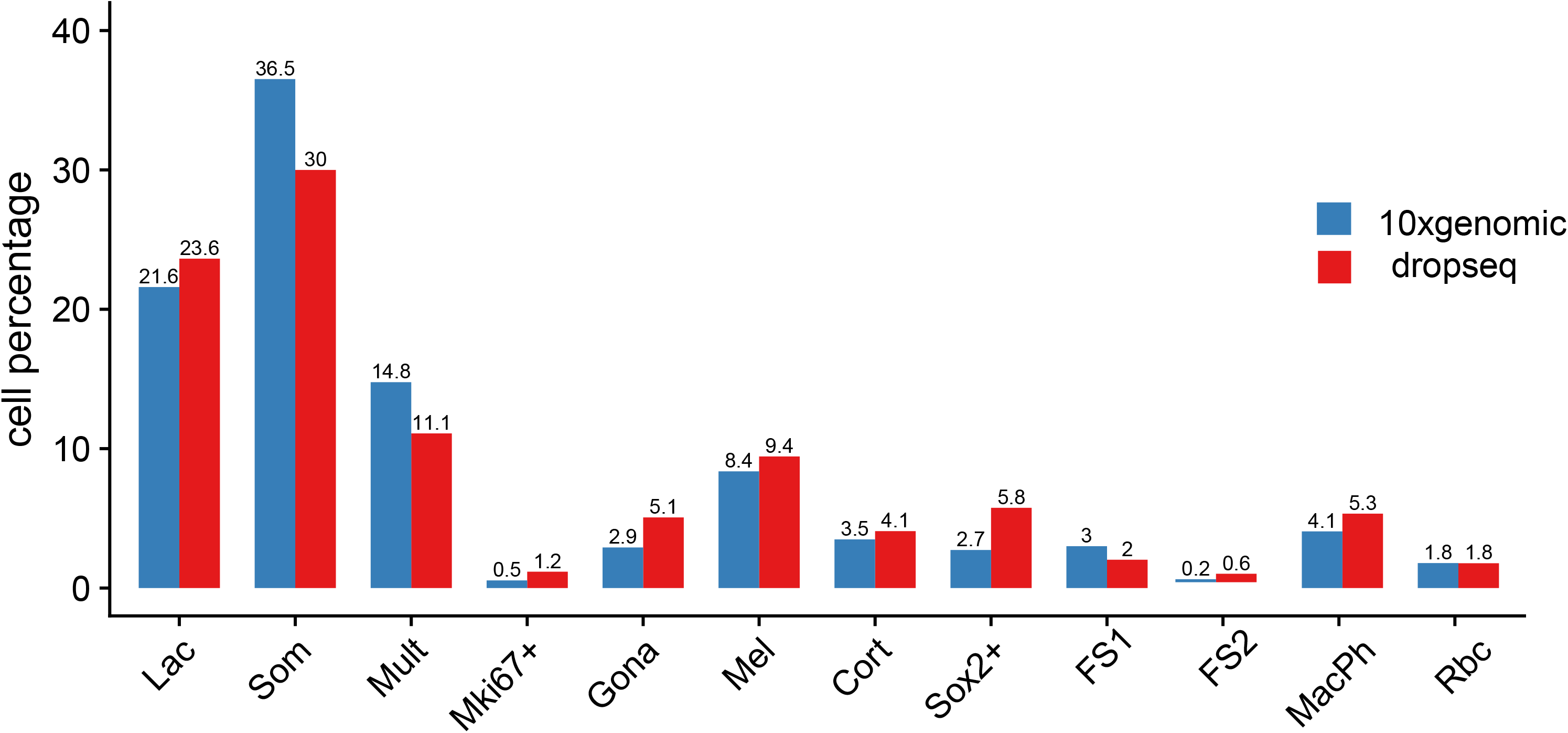
Single cell RNA-seq analysis of 3,927 cells using the 10X Genomics platform. The bar plots compare the cell distribution of cells across each types using the Drop-seq and 10X genomic technologies.

**Figure S4.**
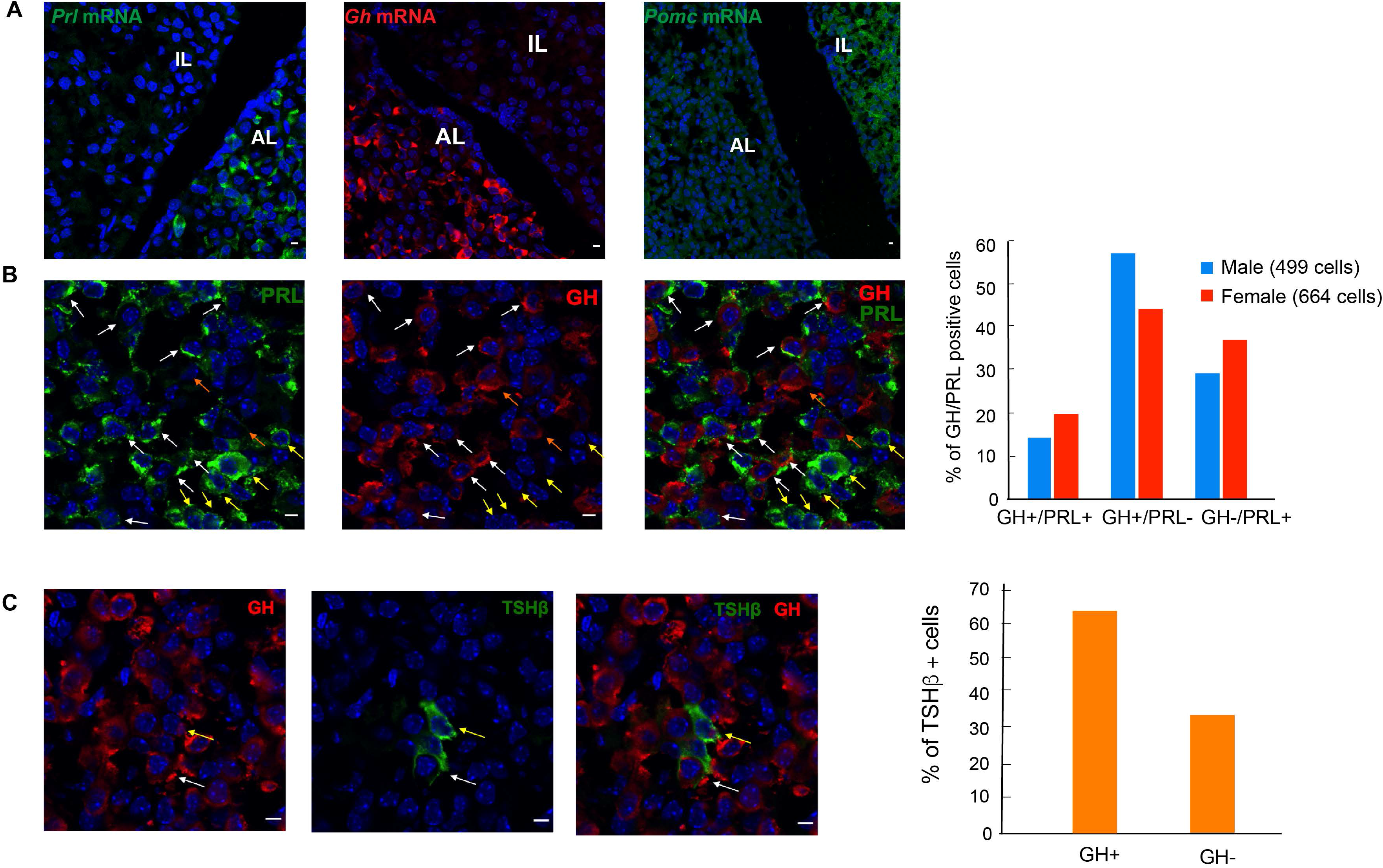
RNA FISH and IF analyses in adult pituitary tissue sections detect the presence of the multi-hormone cells at both mRNA and protein levels. **A. RNA FISH demonstrates that the morphology of the adult pituitary is conserved in pituitary tissue sections.** This analysis was performed on pituitary tissue sections of 8-week old male and female mice. **Left panel:** RNA FISH shows that the robust expression of the *Prl* mRNA (green) is restricted to the anterior lobe (AL). **Middle panel:** RNA FISH shows the robust expression of the *Gh* mRNA (red) is restricted to the anterior lobe (AL). **Right panel:** RNA FISH shows the broad expression of *Pomc* mRNA (grey) in the intermediate lobe (IL) and anterior lobe (AL). **B. IF analysis performed on 8-week old female mice confirms co-expression of GH and PRL proteins in pituitary cells.** The IF was performed in the pituitary tissue sections of an 8-week old female mouse using antibodies specifically for PRL and GH. **Left panel:** IF detection of PRL protein (green). Scale bar = 5 µm. **Middle panel:** IF detection of GH protein (red) in the same field as in left panel. Scale bar = 5 µm. **Right panel:** The merged images of the PRL IF and GH IF. The white arrows indicate the cells with robust levels of GH protein and PRL protein. The yellow arrows indicate cells with robust level of PRL but without GH protein signal. The orange arrows indicate cells with robust level of GH protein but without PRL protein signal. **Histogram:** co-expression of GH and PRL in the pituitary of male or female mouse. The IF analysis was performed in pituitary tissue sections of one male mouse (499 GH and/or PRL expressing cells were analyzed) and one female mouse (664 GH and/or PRL expressing cells were analyzed). **C. IF analysis identifies cells co-expressing of TSHβ and GH proteins *in vivo*.** IF was performed in the pituitary tissue sections of an 8-week old male mouse using antibodies specifically for GH (red in the left panel) and TSHβ (green in the middle panel). The right panel is the merged images of the IF. Two TSHβ positive cells are indicated with arrows. One of these cells is also expressing high level GH (white arrow) while the other cell is expressing relative lower level of GH (yellow arrow). **Histogram: Quantification of cells co-expressing TSHβ and GH proteins.** Of 64 TSH(+) cells detected in this study, 39 were also positive for GH (61%).

**Table S1. List of marker genes in each cluster.**

**Table S2. The cell numbers in each cluster and each sample.**

